# Hypoxia-inducible factors drive miRNA-mediated downregulation of *L2HGDH* and *HIF1A* in clear cell renal cancer independent of 14q deletion

**DOI:** 10.1101/2024.12.01.625494

**Authors:** Soma Seal, Rebecca A. Luchtel, Talia Akram, Bogdan Isaila, Kent T. Perry, Daniele Procissi, Sassan Hafizi, Niraj K. Shenoy

## Abstract

Clear cell renal cell carcinoma (ccRCC) is characterized by pseudohypoxic signaling as well as metabolic and epigenetic aberrations of increasing severity during its pathogenesis and progression. Upon identifying that the markedly lower expression of *L2HGDH* and *HIF1A* in ccRCC tumors is not explained only by 14q deletion (⁓40% of ccRCC), we sought to elucidate the mechanisms underscoring their down-regulation independent of 14q deletion, given the marked translational and scientific importance. While lower *L2HGDH* was found to portend a strikingly worse prognosis in a multivariate survival analysis of ccRCC patients (n=509; TCGA) highlighting its translational importance, the lower expression of *HIF1A* in ccRCC despite the upregulated HIF1α protein (secondary to VHL loss) is of significant scientific interest. Using a comprehensive array of in vitro assays with genetic manipulation, clinicogenomic bioinformatic analyses of large ccRCC datasets, and the *Cdh16-*Cre *Vhl*-fl mouse model with kidney specific *Vhl*-loss, we demonstrate that the downregulation of both *L2HGDH* and *HIF1A* in ccRCC is driven by the hypoxia-inducible factors via miR21-3p and miR155-5p, respectively, during pathogenesis and progression. In doing so, we unveil a conserved inhibitory feedback loop for *HIF1A* in ccRCC (HIFs→miR155→*HIF1A*), providing a mechanistic explanation for *HIF1A* downregulation despite elevated HIF1α protein. Furthermore, and most importantly, we show that inhibition of hypoxia-inducible factors reverses the adverse loss of *L2HGDH* and hydroxymethylcytosine in ccRCC, implicating both HIF2α and HIF1α in these metabolic-epigenetic aberrations.

**Translational Statement:** This study demonstrates that hypoxia-inducible factors (HIFs) drive early and sustained metabolic and epigenetic dysregulation in clear cell renal cell carcinoma (ccRCC). Notably, these aberrations begin early in carcinogenesis. We show that inhibition of HIFs can reverse these pathological changes - most significantly, restoring levels of 5-hydroxymethylcytosine, a key epigenetic marker lost in ccRCC due to metabolic inhibition of TET enzymes. Our findings support the integration of HIF2α inhibition earlier in the therapeutic approach for advanced ccRCC. Moreover, the data suggest that dual inhibition of HIF2α and HIF1α - particularly in tumors with high expression of both factors - may yield a more robust reversal of epigenetic abnormalities and merits further investigation.

**Graphical Abstract:** 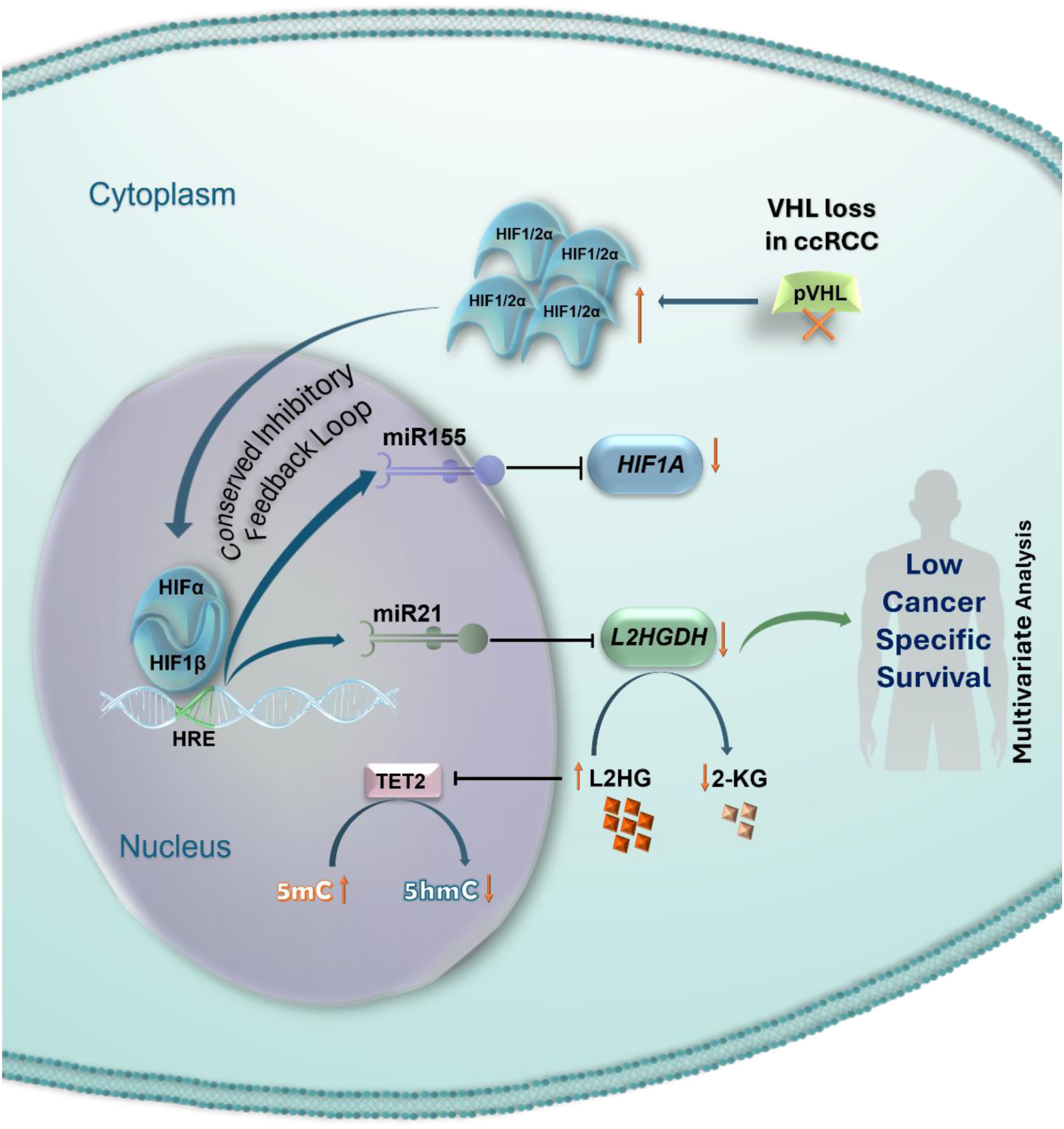

## Introduction

Clear cell renal cell carcinoma is marked by pseudohypoxic signaling, along with progressively severe metabolic and epigenetic abnormalities throughout its development and progression.^1–3^ Around 40% of ccRCC is also characterized by a large 14q deletion, which portends worse prognosis.^4^ *L2HGDH* and *HIF1A* are two of the most well studied genes involved in 14q-deleted ccRCC, with prognostic and scientific significance.^5^

Lower *L2HGDH* expression and consequently lower protein expression has been shown to result in accumulation of oncometabolite L2HG, resulting in competitive inhibition of 2-oxoglutarate dependent dioxygenases including TET enzymes, leading to the adverse loss of hydroxymethylcytosine in ccRCC.^1, 6, 7^

Unlike L2HGDH, however, there is no correlation between *HIF1A* expression and HIF1α protein expression in ccRCC, given the primarily post-translational regulation of HIF1α.^5^ Despite the significantly lower *HIF1A* expression in ccRCC compared to normal kidney, HIF1α protein is markedly upregulated because of biallelic *VHL* loss and reduced/ absent ubiquitination and proteasomal degradation of HIFα isoforms (primarily HIF1α and HIF2α).^8–11^ While HIF2α has been well established as the primary oncogenic isoform in ccRCC, the role of HIF1α in ccRCC has been heavily debated, with the more recent studies indicating a supporting pro-oncogenic role contrary to the prior studies that had claimed a tumor-suppressive role for HIF1α in ccRCC.^4, 5, 12–15^ However, while *HIF2A* is over-expressed, *HIF1A* is under-expressed in ccRCC tumors.

Upon identifying that the markedly lower expression of *L2HGDH* and *HIF1A* in ccRCC tumors compared to normal kidney is not explained only by 14q deletion, we sought to elucidate the mechanisms governing their down-regulation independent of 14q deletion, given the marked translational and scientific importance. We used clinicogenomic bioinformatic analyses from large datasets (TCGA and CPTAC) for screening and hypotheses generation, a comprehensive array of assays with ccRCC and precancerous cell lines with genetic manipulation for hypotheses testing, a mouse model of kidney-specific *VHL* deficiency to validate the molecular pathways *in vivo,* and then determined the translational relevance and applications of the mechanisms in relation to recently approved HIF2α small molecule inhibitor, belzutifan.

## Methods

### Cell Lines

We obtained ccRCC cell lines 786-O, 769-P, and A-704 from the American Type Culture Collection (ATCC) and the RCC4 cell line from Lily Wu (UCLA). We cultured 786-O, 769-P, and RCC4 cells in RPMI-1640 media with 10% fetal bovine serum (FBS) and 1% Penicillin/Streptomycin, while A-704 were maintained in DMEM with 10% FBS and 1% Penicillin/Streptomycin. Additionally, we purchased HK-2 kidney cells from ATCC and received HKC-8 cells from Lorainne Racusen (Johns Hopkins University). HK-2 was cultured in DMEM supplemented with 10% FBS and 1% Penicillin/Streptomycin and HKC-8 cells were cultured in DMEM with 2.5% FBS and 1% Penicillin/Strptomycin.

### miRNA mimic and inhibitor transfection

Mature miRNA mimics for miR21-3p and miR155-5p, as well as a negative control (Ambion/Thermo Fisher Scientific), were transfected into HK-2 and HKC-8 cells using Lipofectamine 3000 (Invitrogen) at a final concentration of 30 nM. Similarly, transfected mature miRNA inhibitors for miR21-3p, miR155-5p and negative control (Ambion/Thermo Fisher Scientific) into 786-O cells at a final concentration of 150 nM using Lipofectamine RNAiMAX (Invitrogen). All transfections were conducted in triplicate and verified by qPCR for miR21-3p and miR155-5p.

### siRNA transfection

siRNA targeting *HIF1A*, *EPAS1*, and non-targeting control were purchased from Horizon Discovery (Dharmacon ON-TARGETplus SMARTpool siRNA). Each siRNA was transfected at a concentration of 50 nM using RNAiMAX. Cells were harvested at 48-h post-transfection for validation of knockdown and assessment of target genes by PCR.

### Belzutifan treatment

A-704 and 786-O cells were seeded at a density of 1.5×10^5^ and 0.8×10^5^ cells per well of 6 well plate, respectively. Cells were treated with 0-20 µM Belzutifan (MedChemExpress, HY-125840) for 12-24 hours.

### DMOG treatment

Dimethyloxalylglycine (DMOG; Cayman Chemical) was solubilized in PBS and prepared fresh for each experiment. HCK-8 cells were treated with 1 mM DMOG and harvested at 24 hours for miRNA and 48 hours for mRNA and protein analysis.

### HIF2α Overexpression

786-O and HKC-8 cells were transfected with 2.5µg of pcDNA3.1-HA (Addgene plasmid #128034; gift from Oskar Laur) and HA-*EPAS1*/HIF2A-pcDNA3 (Addgene plasmid #18950; gift from William Kaelin^16^) expression vectors using Lipofectamine 3000 (Invitrogen) following the manufacturer’s protocol. Cells were harvested 24- or 48-hours post-transfection for RNA or protein analysis.

### Quantitative Real-Time PCR

Total RNA was extracted from using the miRNeasy kit (Qiagen) following the manufacturer’s instructions. The cDNA was then synthesized using I-Script (Biorad) and TaqMan MicroRNA Reverse Transcription Kit (Thermo Fisher Scientific) for mRNA and miRNA analyses, respectively. For small RNA analysis, Taqman primer probe assays were used for miR21-3p (human [hsa-miR-21-3p], 002438; mouse [mmu-mir-21-3p], 002493), miR155-5p (human [hsa-miR155-5p], 467534; mouse [mmu-mir155-5p], 002571), RNU6B (human, 001093), and U6 snRNA (mouse and human, 001973) (Thermo Fisher Scientific). For mRNAs, the primer sequences are listed in Table 1. qPCR was carried out for small RNAs using Taq-Man Universal Master Mix (Thermo Fisher Scientific) and for mRNAs using SSO advanced SYBR green Mastermix (Biorad) following the manufacturer’s instructions. The relative expression of each gene was determined using ΔΔCt normalized to RNU6B or U6 for miRNA and *ACTB* or *GAPDH* for mRNA genes. For use of more than one housekeeper gene for normalization, the geometric mean of housekeeper gene ΔCt values was used in the ΔΔCt calculation. PCR was performed on a Biorad Duet qPCR system.

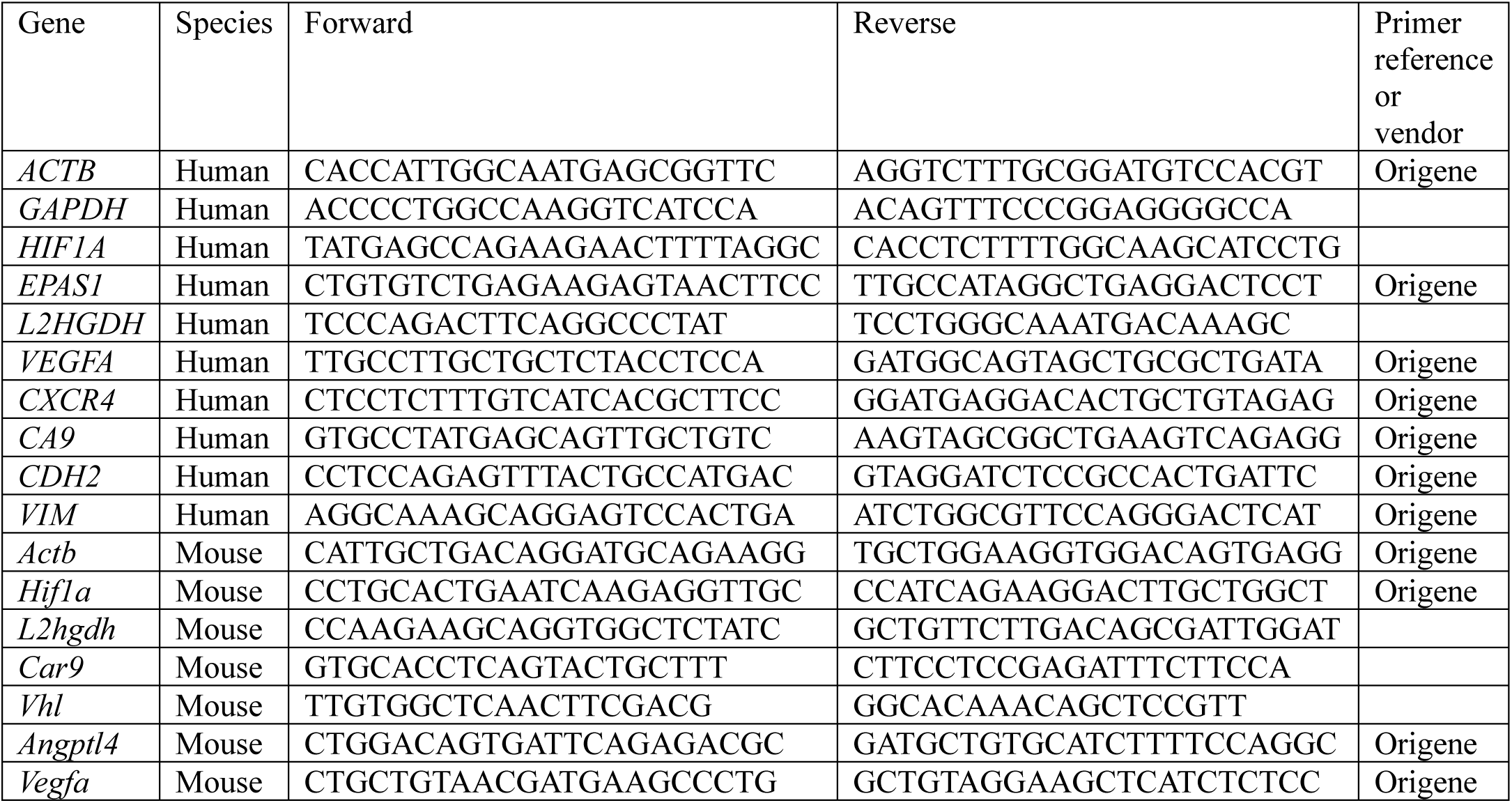

### Immunoblotting

Protein lysates were extracted and subjected to electrophoresis on 4 to 15% Tris·HCl poly-acrylamide gels. The proteins were then transferred to nitrocellulose membranes and blocked using a mixture of Intercept PBS Blocking Buffer (LI-COR) and PBS in a 1:1 ratio. Equal loading was determined by Revert total protein stain (LI-COR). Primary antibodies HIF2α (Novus Biologicals NB100-122), L2HGDH (Proteintech 15707-1-AP), HIF1α (clone 54; BD Biosciences or Novus Biologicals NB100-479), and β-Actin (Novus Biologicals, NB600-501; AC-15) were incubated with the membranes. The proteins of interest were detected using IRDye 800CW and 680RD secondary antibodies (LI-COR) on a LI-COR Odyssey XF Imaging System. Quantification was performed using Empiria Studio software. Protein targets were normalized to total protein or β-Actin and expressed as fold change relative to the control.

### Dot Blot Assay for 5-hydroxymethylcytosine Analysis

Genomic DNA from cell lines and mouse kidneys was extracted using Qiagen DNeasy Blood and tissue kit (#69504). Following quantification, DNA was denatured by adding 2 N NaOH/50 mM EDTA) and incubated at 95 °C for 10 mins. Samples were quickly placed on ice followed by neutralization using ice cold 2 M ammonium acetate. Denatured DNA was spotted on a positively charged nylon membrane (Amersham Hybond N+) using a Biorad Bio-Dot apparatus following manufacturer’s protocol. Hybridization was done by baking for 1 hour at 75 °C. The membranes were then blocked in 5% milk/TBST for 1 h followed by primary antibody incubation at 4 °C overnight at dilution of 1:10,000 rabbit anti hmC (catalog no. 39069, Active Motif). The blots were washed with TBST followed by incubation with secondary 1:3,000 anti-rabbit-HRP for 5-hmC for 1 h, washed, and developed using LI-COR ECL substrate followed by imaging in a LI-COR Odessey XF. Methylene blue stain (0.04% in 0.5 M sodium acetate, pH 5.2) was used as loading control. The nylon membrane with methylene blue spots were scanned using LI-COR D-Digit scanner.

### 2-HG Measurement

L-2HG and D-2HG were measured by LC-MS. Cells cultured in 60 mm dishes were washed with ice-cold PBS and treated with 1ml of pre-chilled 100% methanol with 10µl of 500µM ISTD internal standard (ISTD) (2S)-2-Hydroxyglutaric Acid Disodium Salt-d5 (LGC standards). The plates were incubated at -80 °C for 15 minutes before scraping into microcentrifuge tubes. The resulting cell lysate was centrifuged at 15,000 x g at 4 °C for 15 minutes to pellet insoluble material, and the supernatant was separated into new microcentrifuge tubes for drying. Standards were prepared in 158µl of 100% methanol by adding 10µl of 20 µM L-2HG (L-α-Hydroxyglutaric acid disodium salt (≥98%), catalog # 90790, Sigma) or 10µl of 20 µM D-2HG (D-α-Hydroxyglutaric acid disodium salt, catalog # 16859, Sigma) separately, followed by 2 µl of the 10 mM sodium lactate, 2µl of 500 µM ISTD and 28µl of LC-MS water.

Metabolite extracts were transferred into microcentrifuge tubes and dried using a Speedvac roto-evaporator to remove all solvents. Subsequently, the derivatization reagent was prepared by dissolving DATAN (diacetyl-L-tartaric anhydride (DATAN), catalog # AC336040050, Fisher Scientific) in acetonitrile : acetic acid (4:1, v/v) to a concentration of 50 mg/mL. To initiate derivatization, 50 µL of the DATAN solution was added to the dried samples, vortexed to ensure thorough mixing and then incubated at 72°C for 2 hours. After incubation, samples were allowed to cool to room temperature and briefly centrifuged. The derivatized samples are then diluted with 50 µL of acetonitrile : acetic acid (4:1, v/v), vortexed, spun briefly, and transferred to a clean microcentrifuge tube for LC-MS analysis. For each sample, peak area for each metabolite is divided by the ISTD peak and normalized to total protein. Relative abundance of each metabolite was compared between groups and reported as fold change.

### 3-UTR Reporter Assay

*L2HGDH* (HmiT019408-MT06) and *HIF1A* (HmiT149031-MT06) miTarget 3′UTR miRNA reporter plasmids and pEZX-MT06 control vector were purchased from Genecopoeia. HKC-8 cells were co-transfected with 100 nM miRNA mimics and 100 ng reporter constructs using Lipofectamine 3000 in a 96-well plate. After 24h incubation, luciferase activity was measured using the Luc-Pair Duo-Luciferase assay kit 2.0 procured from Genecopoeia (#LF001) on a Biotek Cytation 3 plate reader following the manufacturer’s instructions.

### Chromatin Immunoprecipitation (ChIP)

ChIP was performed on untreated cell lines using the SimpleChIP Enzymatic Chromatin IP Kit (Cell Signaling). Chromatin was sheared using a combination of MNase digestion and ultrasonication (Covaris) to yield predominantly mono- and di-nucleosomes for each cell line. Approximately 4×10^6^ cells were used per IP with 3.65 µg of antibodies targeting HIF2α (Novus Biologicals, NB100122) or normal rabbit IgG (Cell Signaling) antibodies. Quantitative real-time PCR was used to assess enrichment using the following primer sets: *MIR155* locus (F), CGCGGTGCAGGAAAGTACA; *MIR155* locus (R), CCCCTTTCCCTTTCTCGTGG; *MIR21* ∼55 kb upstream (F), GAAACCAGCTTCCCTGTAACCA; *MIR21* ∼55 kb upstream (R), AACCCTCTAGAGTCCACGCTG. PCR was performed on a Biorad Duet qPCR system using SsoAdvanced Universal SYBR® green Mastermix (Biorad). Enrichment was expressed as percentage of signal relative to Input.

### Chromatin Immunoprecipitation (ChIP)-seq analysis

HIF2α and HIF1β ChIP-seq data in 786-O cells were obtained from GSE34871^17^ and GSE67237^18^. HIF1α, HIF2α, and HIF1β ChIP-seq data in RCC4 cells were obtained from GSE120885^19^. ChIP-seq data were visualized using IGV. DNA sequences from peaks corresponding to *MIR21* and *MIR155* loci were obtained using UCSC genome browser. Hypoxia response elements (HREs) were identified within these peaks for confirmatory ChIP, and peak coordinates were used for enhancer annotation.

### Enhancer analysis

The Human Active Enhancer to interpret Regulator variants (HACER) database was used to annotate enhancers and target genes within specified genomic coordinates.^20^

### Wound healing assay

RCC4 cells were transfected with *HIF1A* and non-targeting control siRNA. 24 h post-transfection, three separate wounds were made per treatment group. Wounds were imaged at 0, 24, and 48 hours at 4x magnification. ImageJ was used to calculate the percent of initial wound (0 h timepoint) was remaining at 24- and 48-hours.

### shRNA transduction

Lentivirus for *HIF1A* (TRCN0000318677) or non-targeting control in the TRC2 pLKO.1 vector (MISSION Lentiviral shRNA, Sigma) was transduced into RCC4 cells using Lenti-X Packaging Single Shots (VSV-G; Takara). Puromycin was used to select and maintain transduced cells. Single cell colonies were selected to ensure consistent knockdown level. Knockdown was assessed by qPCR.

### Invasion assay

Using stable *HIF1A* or non-target control shRNA-expressing RCC4 cells, 10,000 cells were seeded on each 8 µm Matrigel invasion chamber (Corning BioCoat Matrigel Invasion Chambers). Cells were allowed to invade for 24 hours, and were then processed according to manufacturer’s instructions. Invaded cells were visualized with methylene blue staining and captured at 10x magnification.

### Mouse Study

All animal work was approved by Northwestern University Institutional Animal Care and Use Committee. *Cdh16*-Cre (B6.Cg-Tg(Cdh16-cre)91Igr/J; Jax Stock#012237) and *Vhl*-fl (B6.129S4(C)-Vhltm1Jae/J; Jax Stock# 012933) mice were purchased from Jax Laboratories. These strains were crossed by Northwestern CCM Breeding Service and genotypes were determined by Transnetyx. From this cross, age-matched offspring with Cre^+^*Vhl*^fl/fl^ (n= 4; 3 male, 1 female) and Cre^+^*Vhl*^wt/wt^ (n= 3; 1 male, 2 female) genotypes were maintained until 10 months of age for molecular and phenotypic kidney characterization. All age-matched mice generated were included in the study. MRI imaging was performed on age-matched Cre^+^*Vhl*^fl/fl^ and Cre^+^*Vhl*^wt/wt^ mice at 10 months of age at the Northwestern University Center for Translational Imaging (CTI) (Wt, n=2; KO, n=2). All seven mice were then sacrificed at 10 months of age and kidneys were harvested for protein, DNA, and RNA extraction. Data for each mouse are reported individually. (We have used the ARRIVE reporting guidelines^21^)

### Bioinformatic analyses

The UCSC Xena^22^ and Cbioportal^23^ platforms were used to access TCGA KIRC data for Figs. 1A, 1B, S1A, S1B, 2C, 2D, 3E, 3F. (Analysis was done using Microsoft Excel for these figures. Heatmapper.org was used to generate heatmaps for Figs 2C, 2D, 3Ei, 3Fi.)

**Figure 1.**
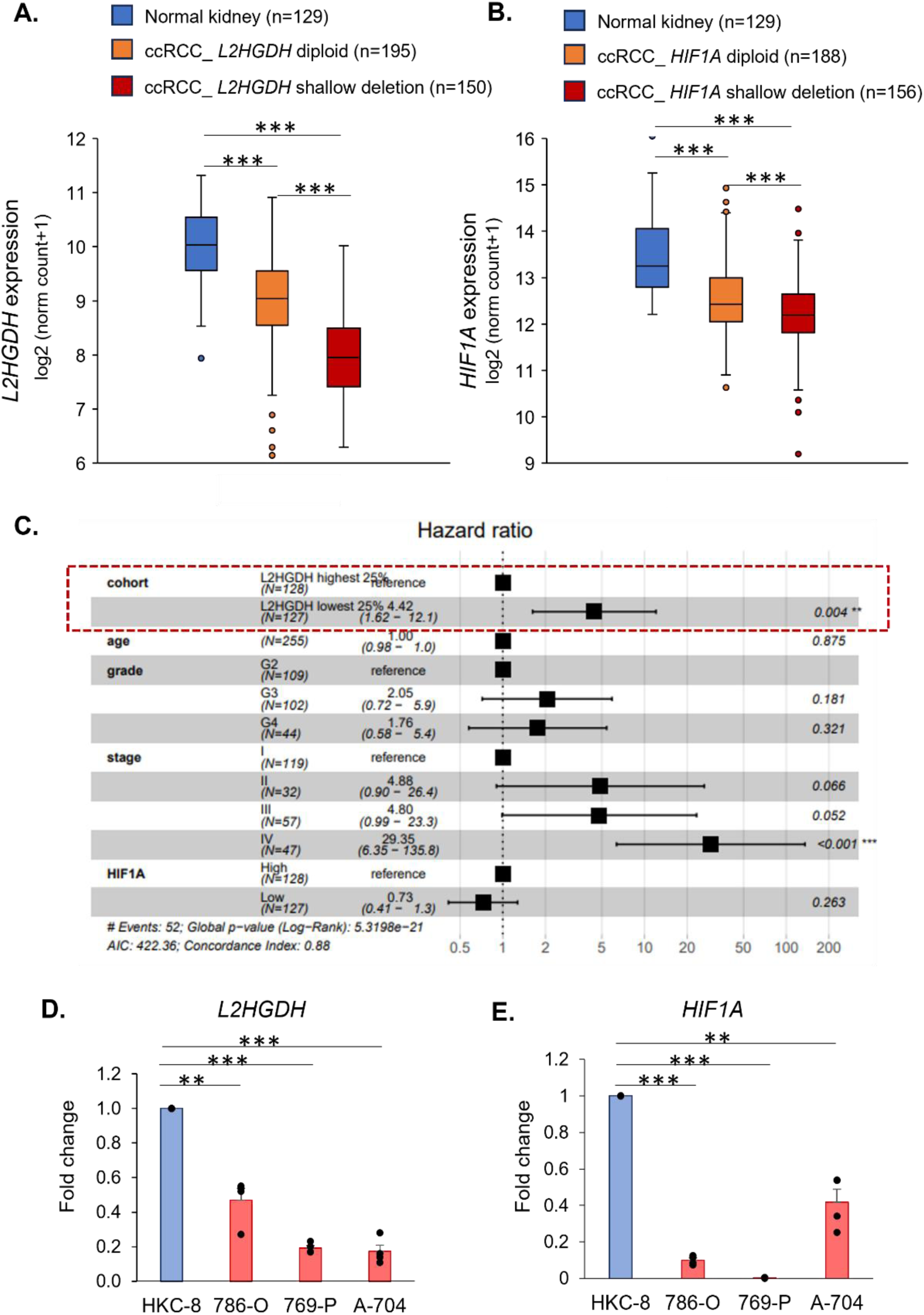
*L2HGDH* and *HIF1A* are markedly downregulated in ccRCC even without 14q deletion, with loss of *L2HGDH* portending markedly worse survival in multivariate analysis. **A, B.** TCGA KIRC data revealing significant downregulation of both *L2HGDH* **(A)** and *HIF1A* **(B)** even in 14q diploid ccRCC compared to normal kidney, indicating mechanism/s of downregulation other than 14q loss. [***, p<0.001; 2-sided t-test] (Data for paired adjacent normal kidney versus ccRCC tumor is shown in Supplement, also showing significant downregulation of *L2HGDH* and *HIF1A*). **C.** Lower *L2HGDH* expression is associated with strikingly lesser cancer-specific survival in ccRCC in a multivariate analysis. After adjusting for grade, stage, age, and *HIF1A* expression, the lowest quartile of *L2HGDH* expression was associated with a 342% increase in hazard of death from ccRCC compared to the uppermost quartile (HR 4.42; 95% CI: 1.62-12.1; p=0.004; dotted red line box), underscoring the critical need for delineating mechanisms of its downregulation. **D, E.** ccRCC cell lines 786-O, 769-P, A-704, have markedly lesser expression of *L2HGDH* **(D)** and *HIF1A* (**E**) compared to normal kidney cell line HKC-8. (n=4; mean+/-SE) [**, p<0.01; ***, p<0.001; paired t-test].

**Figure 2.**
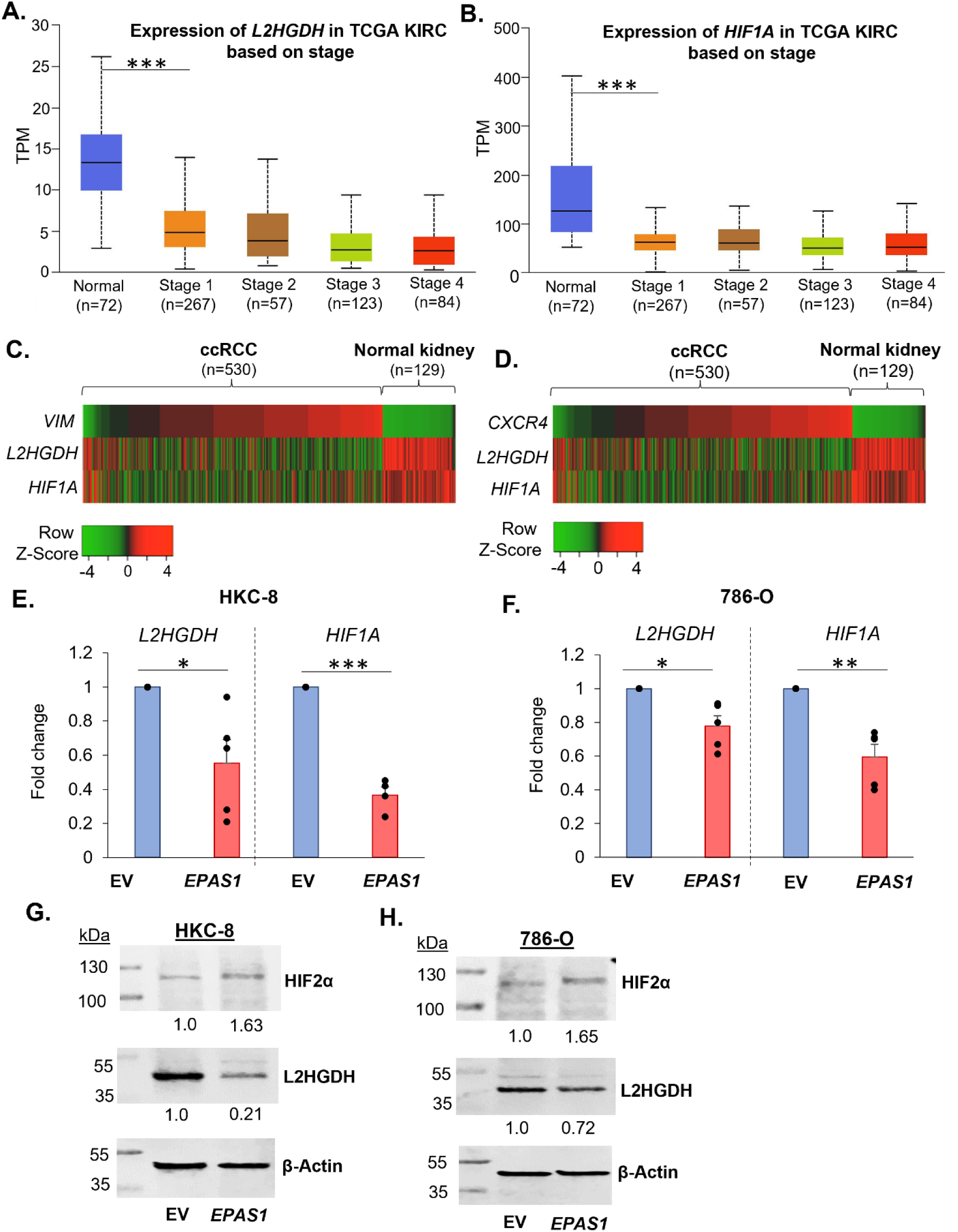
HIF2α downregulates *L2HGDH* and *HIF1A* in ccRCC early in its progression. **A, B**. Loss of *L2HGDH* (**A**) and *HIF1A* (**B**) expression occurs early in ccRCC progression, with a marked down-regulation already in stage 1 compared to normal kidney. Data from TCGA KIRC; UALCAN interface (***, p<0.001). **C, D**. Heatmap showing HIF2α target genes *VIM* (**C**) and *CXCR4* (**D**), having a strong negative correlation with *L2HGDH* (R= -0.59 and -0.57, respectively; p<0.001 for each) and with *HIF1A* (R= -0.37 and -0.35 respectively; p<0.001 for each). Data derived from TCGA KIRC. **E, F.** qRT-PCR revealing that transfection of *EPAS1* (encoding HIF2α) plasmid in HKC-8 (**E**) led to marked downregulation of *L2HGDH* and *HIF1A* (n=5, mean+/- SE). Transfection of *EPAS1* in 786-O (**F**) also led to down-regulation of *L2HGDH* and *HIF1A* (n=5, mean+/- SE) (*, p<0.05; **, p<0.01; ***, p<0.001; paired t-test). **G, H**. Representative Western blots showing downregulation of L2HGDH with transfection of *EPAS1* plasmid in HKC-8 (**G**) and in 786-O (**H**), with a much greater degree of downregulation in HKC-8, consistent with qRT-PCR findings. Quantification shown relative to total protein expression.

**Figure 3.**
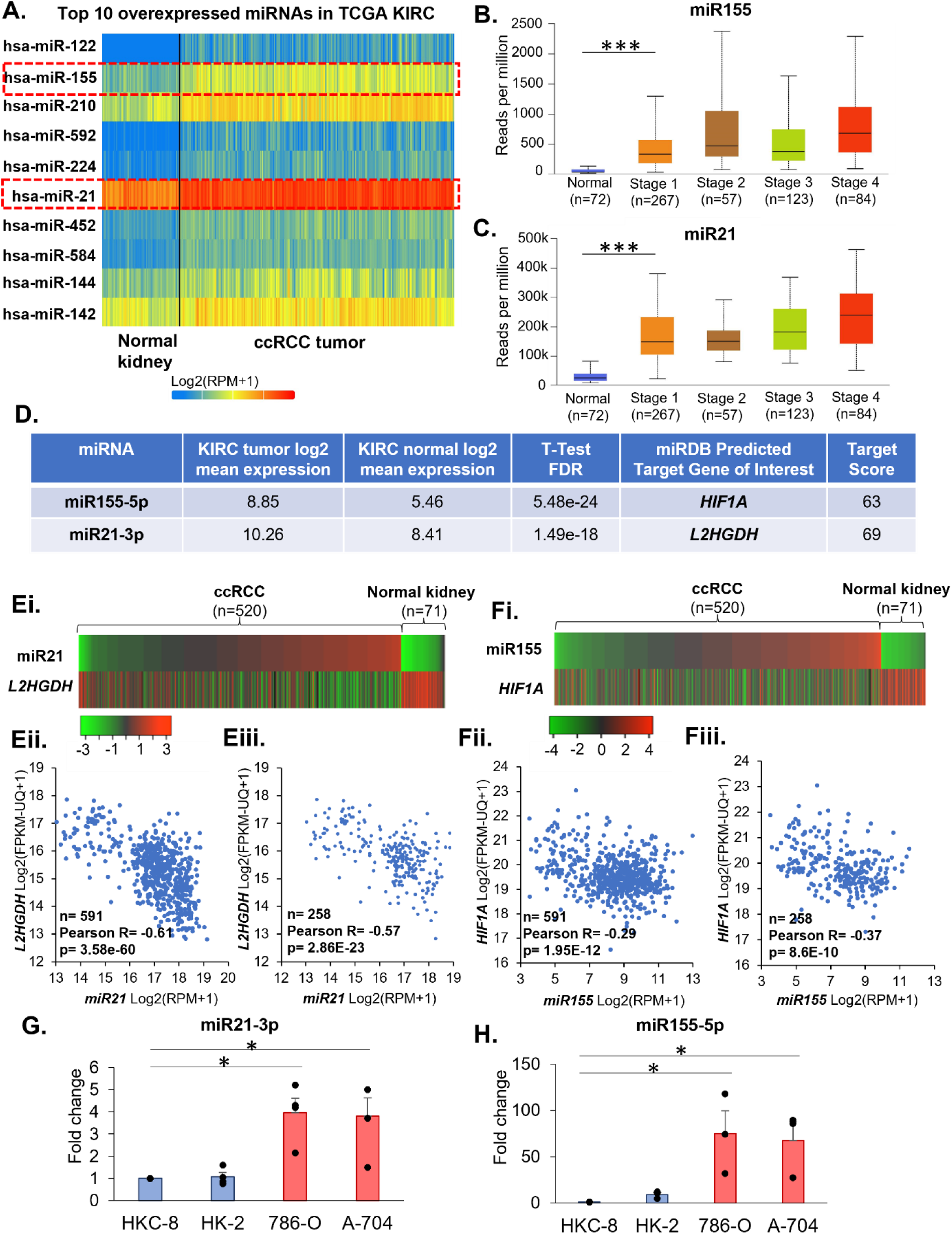
miR21 and miR155 are markedly upregulated early in ccRCC progression and are predicted to target *L2HGDH* and *HIF1A*, respectively, with strong inverse correlation. **A**. Heatmap showing the top ten over-expressed miRNAs in ccRCC (TCGA KIRC). miR155 is the 2^nd^ most upregulated miRNA and miR21 is the 6^th^ most upregulated miRNA in ccRCC. **B, C.** Both miR155 and miR21 are markedly upregulated early in ccRCC progression (TCGA KIRC). **D**. miR21-3p is predicted to directly inhibit *L2HGDH* with a target score of 69; miR155-5p is predicted to directly inhibit *HIF1A* with a target score of 63 (miRDB). miR21-3p and miR155-5p are significantly upregulated in ccRCC tumor compared to normal kidney (log2 mean expression and T-test FDR shown; TCGA KIRC; OncomiR) **Ei-iii, Fi-iii**. Heatmap and scatter plots showing a marked negative correlation between expression of miR21 and *L2HGDH* (**Ei-Eiii**) and that between miR155 and *HIF1A* (**Fi-Fiii**) in normal kidney and ccRCC (TCGA KIRC). (**Ei-ii**) and (**Fi-ii**) include normal kidney and ccRCC from TCGA KIRC, whereas (**Eiii**) and (**Fiii**) include normal kidney and 14q diploid ccRCC only (annotation extracted from cbioportal), demonstrating that the strong negative correlation exists regardless of 14q deletion. **G, H.** qRT-PCR showing a markedly higher expression of miR21-3p and miR155-5p in ccRCC cell lines compared to normal kidney cell lines HKC-8 and HK-2 (n=3, error bars represent SE).

The CVCDAP^24^ platform was used to perform multivariate survival analyses with TCGA KIRC data shown in Fig. 1C, S1C.

The UALCAN^25^ platform was used for analysis of ccRCC samples from TCGA KIRC (Figs 2A, 2B, 3A, 3B, 3C, S1D, S1E, S3A, S3B, S6A-D).

The UCSC Xena platform was used to generate Fig S3C, S3D.

IGV (Integrative Genomic Viewer)^26^ was used to visualize ChIP-seq data, in Figs 5C, 5D, 8A, 8B, S5A, S5B.

The UCSC Genome browser^27^ was used to obtain DNA sequences from peaks corresponding to *MIR21* and *MIR155* loci, and to identify hypoxia response elements (HREs) within these peaks.

The miRDB^28^ platform was used for miRNA target gene prediction and the OncomiR^29^ platform was used to access and analyze TCGA KIRC miRNA data-shown in Fig 3D.

GSEA was used for Pathway Enrichment Analysis. ccRCC KIRC data were downloaded from TCGA. Methylation beta values for the available 215 ccRCC cases were downloaded from TCGA-KIRC. A ranked dataset comparing two groups (*CA9* lower versus *CA9* higher or CGI methylation higher versus lower was generated by calculating the product of the fold-change sign and −log(*P* value) for each gene. This ranked file was uploaded to GSEA for analysis of pathways that were enriched for these comparisons.

### Statistical Analysis

The student’s t test was used for statistical analyses (paired/ one-sided/ two-sided indicated in individual figure legends). Statistical significance was considered P < 0.05.

## Results

### *L2HGDH* and *HIF1A* are markedly downregulated in ccRCC even without 14q deletion, with loss of *L2HGDH* portending markedly worse survival in multivariate analysis

Analysis of the TCGA KIRC data revealed significant downregulation of both *L2HGDH* (**Figure 1A**) and *HIF1A* (**Figure 1B**) even in 14q diploid ccRCC compared to normal kidney, indicating mechanism/s of downregulation other than 14q loss. (Data for paired adjacent normal kidney versus ccRCC tumor is shown in **Figure S1A, S1B,** also showing significant downregulation of *L2HGDH* and *HIF1A.*)

We and others have previously demonstrated that lower L2HGDH results in accumulation of oncometabolite L2HG, a competitive inhibitor of 2-oxoglutarate dependent dioxygenases including TET enzymes, leading to adverse loss of hydroxymethylcytosine in ccRCC.^1, 6, 7^

In order to determine the degree to which lower expression of *L2HGDH* is associated with worse outcomes in ccRCC, a Cox regression multivariate survival analysis was performed with the TCGA KIRC dataset, adjusting for confounding variables (**Figure 1C**). Lower *L2HGDH* expression was found to be associated with markedly lesser cancer-specific survival in ccRCC in the multivariate analysis. After adjusting for grade, stage, age, and *HIF1A* expression, the lowermost quartile of *L2HGDH* expression had a 342% increase in the hazard of death from ccRCC compared to the uppermost quartile (HR 4.42; 95% CI: 1.62-12.1; p=0.004, **Figure 1C**; also see **Figure S1C**), underscoring the critical need for delineating mechanisms of its downregulation.

Consistent with what is observed in ccRCC tumors, ccRCC cell lines 786-O, 769-P, A704, had a markedly lower expression of *L2HGDH* and *HIF1A* compared to normal kidney cell line HKC-8 (**Figure 1D, 1E**).

### HIF-2 downregulates *L2HGDH* and *HIF1A* in ccRCC early in its progression

Loss of *L2HGDH* and *HIF1A* occurs early in kidney cancer progression, with a marked downregulation even in stage 1 (**Figure 2A, 2B**) and grade 1 (**Figure S1D, S1E**) disease compared to normal kidney. However, while L2HGDH protein is markedly downregulated early in ccRCC progression compared to normal kidney (**Figure S2A, S2C**), HIF1α is markedly upregulated early in its progression (**Figure S2B, S2D**).

This led to the hypothesis that the early downregulation of *L2HGDH* and *HIF1A* may be directly or indirectly caused by the pathognomonic feature of ccRCC i.e., biallelic inactivation of VHL and stabilization/ increased activity of HIF2α. Further supporting this hypothesis was that known HIF2 target genes such as *VIM*, *CXCR4*, *IGFBP3* were found to have a strong negative correlation with *L2HGDH* (R= -0.59, -0.57, -0.51 respectively; p<0.0001 for each, **Figure 2C, 2D**) and with *HIF1A* (R= -0.37, -0.35, -0.39 respectively; p<0.0001 for each, **Figure 2C, 2D**) with analysis of TCGA KIRC (*IGFBP3* heatmap not included).

To test the hypothesis, we transfected *EPAS1* (gene encoding HIF2α) plasmid in normal kidney cell line HKC-8 and ccRCC cell line 786-O. As hypothesized, transfection of *EPAS1* plasmid in HKC-8 led to marked downregulation of *L2HGDH* and *HIF1A* (**Figure 2E**). *EPAS1* plasmid transfection in 786-O also led to downregulation of *L2HGDH* and *HIF1A* (**Figure 2F**), albeit to a lesser magnitude compared to HKC-8, probably given the higher expression of HIF2α and lower expression of L2HGDH at baseline in 786-O.

Similarly, there was marked L2HGDH protein downregulation with *EPAS1* plasmid transfection in HKC-8 (**Figure 2G**), and to a lesser extent in 786-O (**Figure 2H**); as well as HIF1α downregulation with *EPAS1* plasmid transfection in HKC-8 (**Figure S2E**).

### miR21 and miR155 are markedly upregulated early in ccRCC progression and are predicted to target *L2HGDH* and *HIF1A,* respectively, with strong inverse correlation

We then sought to identify the mechanisms by which HIF2α downregulates *L2HGDH* and *HIF1A*.

Analysis of HIF2α DNA binding by ChIP-seq in 786-O failed to identify binding in the promoter regions of *L2HGDH* and *HIF1A*. We then hypothesized that the downregulation may be miRNA-mediated, similar to HIF2α induced miR210-mediated downregulation of *SDHD* in ccRCC, that we have previously reported.^30^

We identified the top ten over-expressed miRNAs in ccRCC (TCGA KIRC, **Figure 3A**), recognizing that most of these miRNAs would likely be driven by HIFs, directly or indirectly. Out of these miRNAs, we sought to identify miRNAs that were predicted to directly inhibit *L2HGDH* and *HIF1A*. miR21-3p was predicted to directly inhibit *L2HGDH* with a target score of 69; miR155-5p was predicted to directly inhibit *HIF1A* with a target score of 63 (**Figure 3D**). miR155 was the 2^nd^ most upregulated miRNA and miR21 was the 6^th^ most upregulated miRNA in ccRCC (**Figure 3A**), with both miRNAs upregulated early in ccRCC progression (TCGA KIRC) (**Figures 3B, 3C, S3A, S3B**). Higher miR21 and higher miR155 expression were associated with significantly worse survival in ccRCC (TCGA KIRC) **(Figure S3C, S3D)**.

We then mapped miR21 with *L2HGDH* expression, and miR155 with *HIF1A* expression in ccRCC (n = 520) and normal kidney (n = 71) (TCGA KIRC, **Figures 3Ei, Fi**). miR21 expression had a marked negative correlation with *L2HGDH* expression (R = −0.61, P = 3.5e-60, **Figure 3Eii**); and miR155 had a significant negative correlation with *HIF1A* (R = −0.29, P = 1.9E-12, **Figure 3Fii**). In normal kidney and 14q-intact ccRCC, the R values were -0.57 and -0.37 respectively (**Figures 3Eiii, Fiii**).

We then checked the expression of miR21-3p and miR155-5p in ccRCC and normal kidney cell lines. Consistent with what is observed in ccRCC tumors, ccRCC cell lines 786-O and A-704 had a markedly higher expression of miR21-3p and miR155-5p compared to immortalized kidney cell lines HKC-8 and HK-2 (**Figures 3G, 3H**).

### miR21-3p and miR155-5p directly inhibit *L2HGDH* and *HIF1A,* respectively, in ccRCC

We then sought to determine if miR21-3p and miR155-5p inhibit *L2HGDH* and *HIF1A*, respectively.

Transfection of miR21-3p mimic led to downregulation of *L2HGDH* in both HKC-8 and HK-2. Similarly, transfection of miR155-5p mimic led to downregulation of *HIF1A* in both HKC-8 and HK-2. (**Figures 4A, 4B**) Western blots also showed downregulation of L2HGDH and HIF1α with the respective miRNA mimics. (**Figures 4C, 4D** and **Figure S4**)

**Figure 4.**
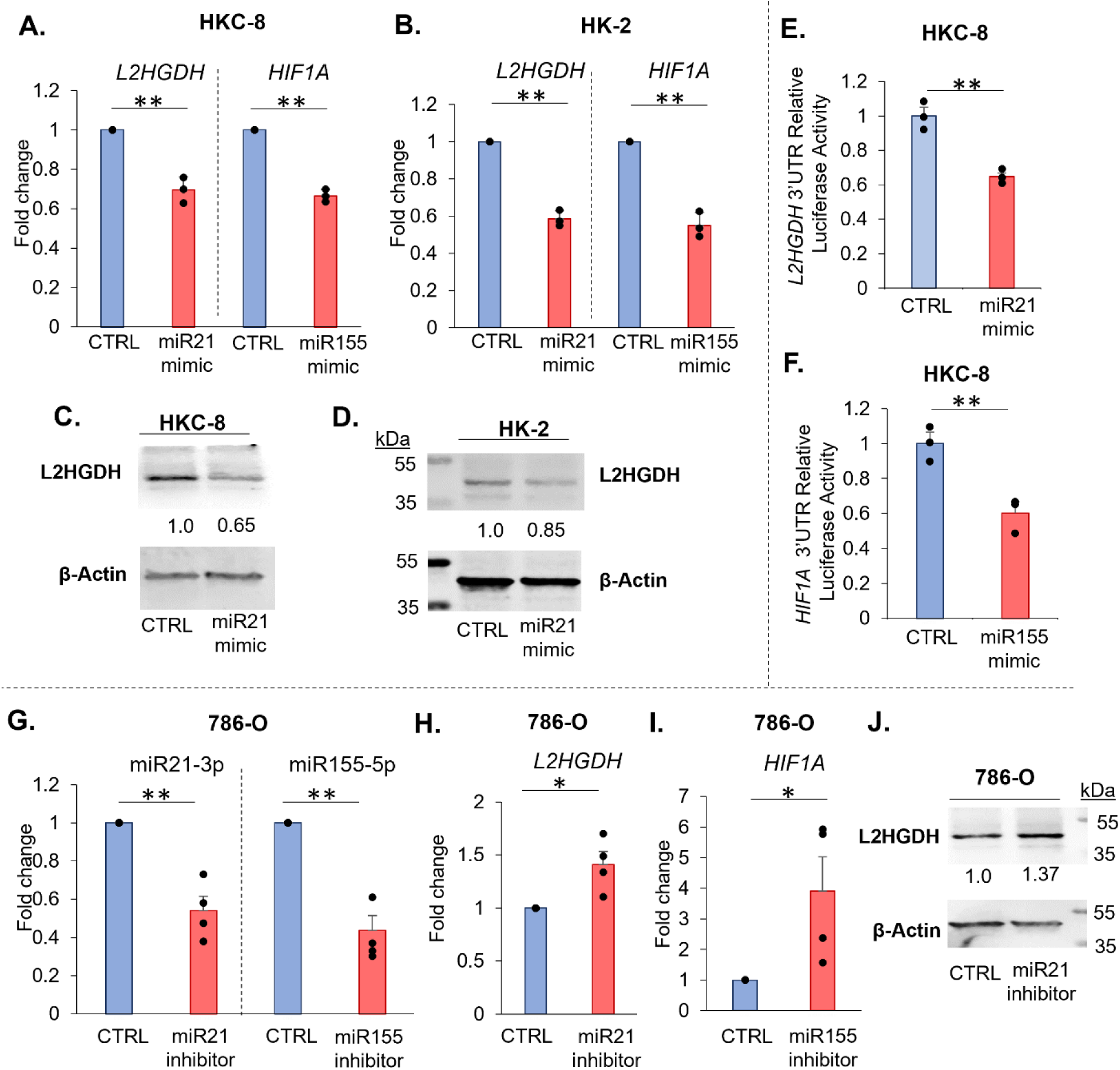
miR21-3p and miR155-5p directly inhibit *L2HGDH* and *HIF1A,* respectively, in ccRCC. **A, B.** Transfection of miR21-3p mimic led to downregulation of *L2HGDH* in both HKC-8 (n=4, P=0.0096) and HK-2 (n=3, P=0.010) cell lines, compared to control miRNA transfection. Similarly, transfection of miR155-5p mimic led to downregulation of *HIF1A* in both HKC-8 (n=4, P=0.0071) and HK-2 (n=3, P=0.0048). (**:P<0.01, paired T-test). **C, D.** Representative Western blots showing downregulation of L2HGDH protein with transfection of miR21-3p mimic in HKC-8 (**C**) and in HK-2 (**D**) normal kidney cells. Quantification shown relative to total protein expression. **E, F.** miR21-3p inhibited *L2HGDH* 3′UTR activity (**E,** P= 0.0028) and miR155-5p inhibited *HIF1A* 3′UTR activity (**F,** P= 0.0081) in HKC-8. (n=3; **:P<0.01; two-sided t-test). **G.** Fold change reduction of miR21-3p and miR155-5p with the inhibitors. **H, I.** qRT-PCR showing upregulation of *L2HGDH* with miR21-3p inhibition (**H,** n=4, P=0.024, paired t-test) and upregulation of *HIF1A* (**I,** n=4, P=0.041, paired t-test) with miR155-5p inhibition in 786-O. **J.** Representative Western blot showing upregulation of L2HGDH with miR21-3p inhibition in 786-O.

In order to determine whether the *L2HGDH* down-regulation by miR21-3p and *HIF1A* down-regulation by miR155-5p in these cell lines was through direct degradation of the transcripts or an indirect mechanism, we co-transfected miR21-3p (or control miRNA) and miR155-5p (or control miRNA) mimics with an *L2HGDH* 3′UTR reporter construct and *HIF1A* 3′UTR reporter construct in HKC-8 cells, and luciferase activity was determined 48 h post-transfection. miR21-3p significantly inhibited *L2HGDH* 3′UTR activity in HKC-8 cells, establishing that miR21-3p directly degrades *L2HGDH* (**Figure 4E**). Similarly, miR155-5p significantly inhibited *HIF1A* 3′UTR activity in HKC-8 cells, establishing that miR155-5p directly degrades *HIF1A* (**Figure 4F**).

Next, we asked if inhibition of miR155-5p and miR21-3p leads to an increase in *HIF1A* and *L2HGDH* respectively in 786-O ccRCC cells. Indeed, transfection of inhibitors of miR21-3p and miR155-5p led to upregulation of *L2HGDH* and *HIF1A*, respectively, in 786-O cells (**Figure 4G-I**). Also, Western blot showed upregulation of L2HGDH protein with miR21-3p inhibition in 786-O (**Figure 4J**).

### miR21-3p and miR155-5p are direct transcriptional targets of HIF2α

The TCGA dataset revealing the markedly higher expression of miR21 and miR155 compared to normal kidney made it likely that they were targets of HIFs. To test this, we experimentally manipulated *HIF2A* expression to determine the role of HIF2α in miR155-5p and miR21-3p expression. In both immortalized kidney (HKC-8) and ccRCC (786-O) cell lines, *EPAS1* transfection significantly increased expression of both miR21-3p and miR155-5p (**Figures 5A, 5B**). To determine whether HIF2α binds to the regulatory regions of *MIR155* and *MIR21*, we visualized HIF2α and HIF1β ChIP-seq data in 786-O (**Figures 5C, 5D**). A binding site was identified at the previously described HRE, ∼ 11kb upstream of *MIR155.*^31^ While binding was not observed within the *MIR21* promoter region, we identified HIF2α binding ∼55kb upstream of *MIR21*, a predicted enhancer for *MIR21* (**Figure 5E**). Binding was also observed in these regions in an independent 786-O ChIP-seq dataset (**Figures S5A, S5B**). To further confirm binding of HIF2α at these sites, we performed HIF2α ChIP in the 786-O and A-704 cell lines, and observed consistent enrichment of HIF2α binding at both *MIR155* and *MIR21* loci (**Figures 5F, 5G, S5C**).

**Figure 5.**
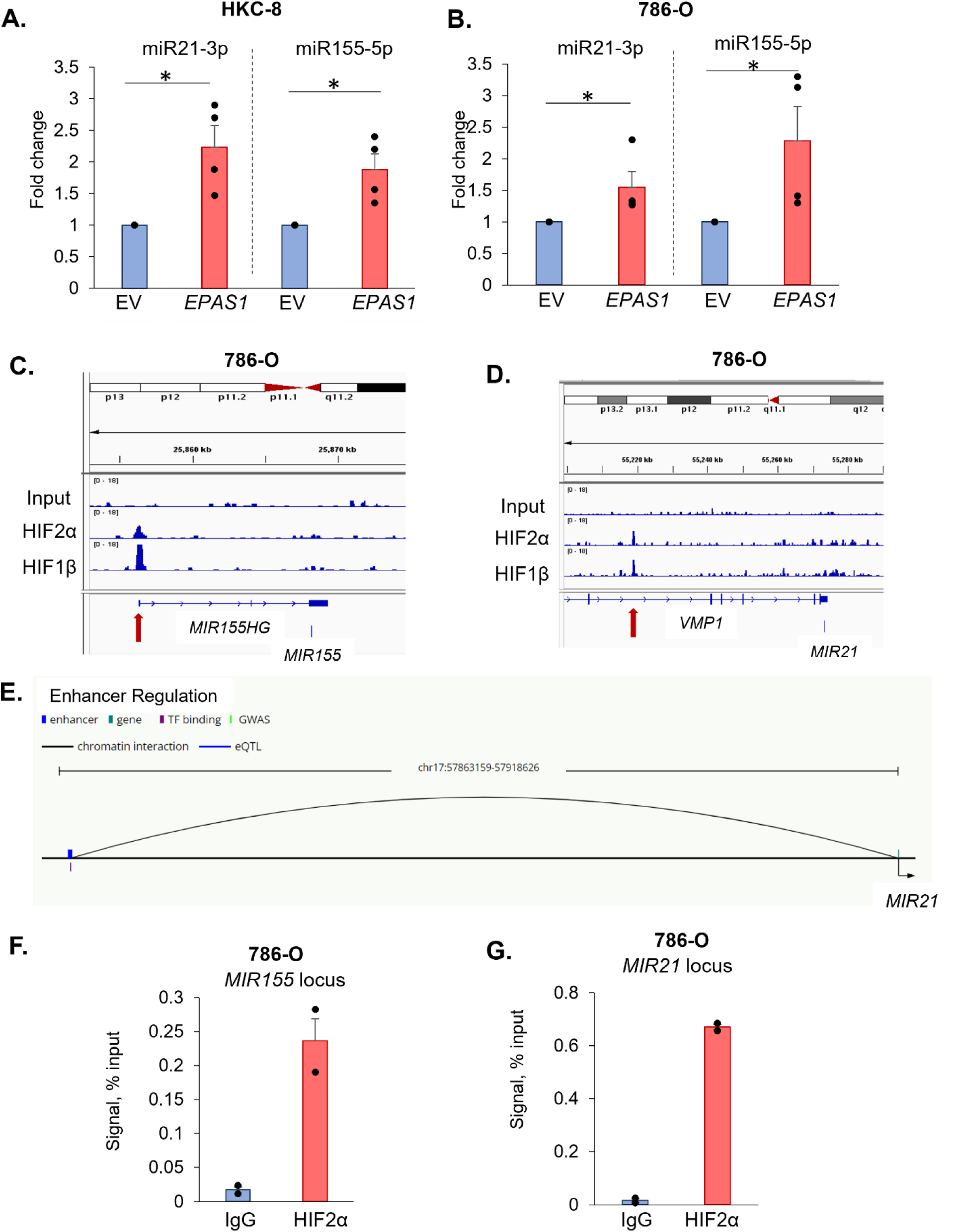
*MIR21* and *MIR155* are direct transcriptional targets of HIF2. **A.** Transfection of *EPAS1* (HIF2A) plasmid in HKC-8 led to upregulation of miR21-3p (n=4, mean+/-SE, p=0.0176) and miR155-5p (n=4, mean+/-SE, P=0.0198). (*:P<0.05, paired T-test) **B.** Transfection of *EPAS1* plasmid in 786-O also led to upregulation of miR21-3p (n=4, mean+/- SE, p=0.0369) and miR155-5p (n=4, mean+/-SE, P=0.048). (*:P<0.05, paired T-test) **C.** ChIP-seq HIF2 (HIF2α, HIF1β) peak identified for *MIR155* in 786-O cells (GSE34871)^17^. [11kb upstream of *MIR155*, overlapping with *MIR155HG* TSS ^31^ predicted as regulatory site for *MIR155* by HIF]. **D.** HIF2 ChIP-seq (HIF2α, HIF1β) peak identified for *MIR21* in 786-O cells (GSE34871), ∼55kb upstream of *MIR21,* but not within the promoter region. **E.** The HIF binding peak ∼55kb upstream of *MIR21* (chr17:57863159-57864676) was identified as an enhancer and shown to interact with the *MIR21* locus (HACER). **F, G**. HIF2α binding to *MIR155* and *MIR21* loci was confirmed by HIF2α ChIP in 786-O.

### *In vivo* validation of the HIF-miRNA-target gene axis in a kidney-specific mouse model of *Vhl* deficiency

To assess whether this relationship was representative of *in vivo* HIF pathway signaling, we generated and utilized the *Cdh16* Cre^+^Vhl^fl/fl^ mouse model with kidney specific *Vhl*-loss. This model is characterized by hydronephrosis formation of increasing severity with age, as confirmed by non-invasive kidney MRI (**Figure 6A**) and consistent with a prior report.^32^ In kidneys harvested from 10-month-old *Cdh16* Cre^+^*Vhl*^wt/wt^ (*Vhl* WT, n=3) and *Cdh16* Cre^+^*Vhl*^fl/fl^ (*Vhl* KO, n=4) mice, we performed gene expression analysis (**Figures 6B, 6C**). We first confirmed *Vhl* knockout and up-regulation of hallmark HIF target genes, *Car9, Angptl4*, and *Vegfa* as confirmation of HIF signaling in *Vhl* KO kidneys (**Figure 6B**). Both HIF1α-specific (*Car9*) and HIF2α-specific (*Vegfa*) target genes were upregulated with *Vhl* knockout. We then assessed miRNA (miR155-5p and miR21-3p) and their target gene expression (*L2hgdh* and *Hif1a*) in this model. Both miR155-5p and miR21-3p were significantly increased in *Vhl* KO kidneys, and this was accompanied by a significant reduction in *L2hgdh* expression, with *Hif1a* expression decreased to a lesser extent (**Figure 6C**). The discord between HIF1α target, *Car9* and *Hif1a* gene expression is consistent with what is found in human ccRCC tumors. Consistent with mRNA expression, L2HGDH protein expression was significantly lower in *Vhl* KO kidneys (**Figure 6D**), together revealing that loss of VHL is sufficient to significantly downregulate L2HGDH. Given the robust decrease in L2HGDH in *Vhl* KO kidneys, we sought to compare DNA hydroxymethylation between *Vhl* WT and *Vhl* KO kidneys. Indeed, we observed a pronounced decrease in 5-hmC in *Vhl* KO kidneys (**Figure 6E**), consistent with the previously described relationship between L2HGDH expression and 5-hmC.^1, 7^

**Figure 6.**
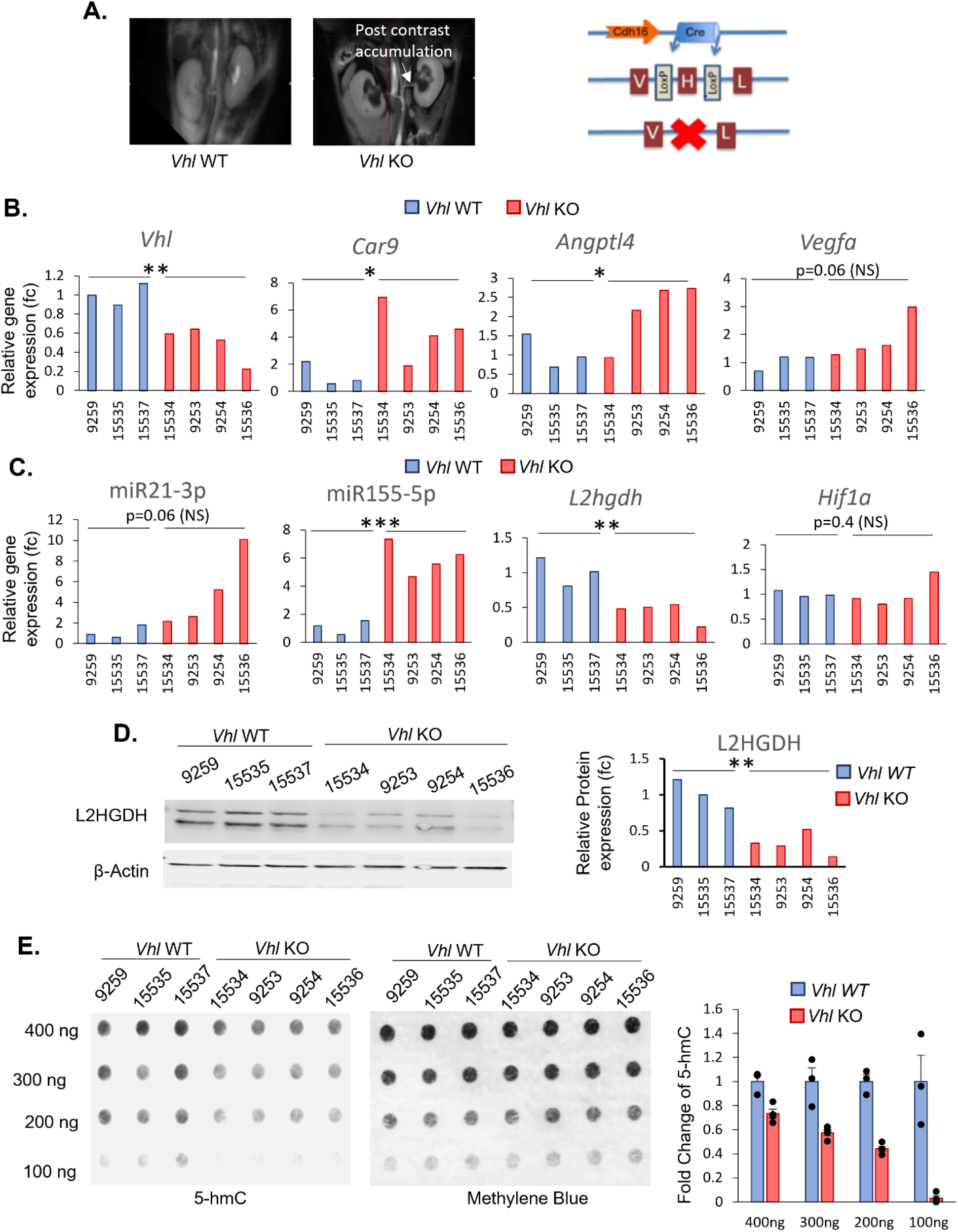
Molecular pathway validation in kidney-specific mouse model of *Vhl* deficiency. **A.** Representative MRI images depicting side-by-side 2D images of *Vhl*^wt/wt^ (Vhl WT) and *Vhl*^fl/fl^ (*Vhl* KO) kidney, enabling visualization of hydronephrosis in *Cdh16* Cre^+^*Vhl^fl^*^/fl^ mice, a penetrant feature in these mice. **B-E.** Kidneys were harvested from n=3 *Vhl*^wt/wt^ (Vhl WT) and n=4 *Vhl*^fl/fl^ (*Vhl* KO) mice at 10 months of age. **B.** Confirmation by qRT-PCR of *Vhl* knockdown (1.00 ± 0.065 vs 0.50 ± 0.094 fc) and upregulation of hypoxia signaling in *Vhl* KO compared to *Vhl* Wt mice (*Car9*, 1.19 ± 0.51 vs 4.37 ± 1.04; *Vegfa*, 1.03 ± 0.16 vs 1.84 ± 0.39; *Angptl4* 1.06 ± 0.26 vs 2.13 ± 0.42 expression fc). Mean ± se. **C.** *Vhl* KO kidneys have upregulation of miR21-3p (1.11 ± 0.36 vs 5.01 ±1.82) and miR155-5p (1.09 ± 0.29 vs 5.96 ± 0.56) and corresponding downregulation of *L2hgdh* (1.01 ± 0.12 vs 0.44 ± 0.074) and *Hif1a* (1.00 ±0.035 vs 1.02 ± 0.15) gene expression fc. Mean ± se. **D.** Downregulation of L2HGDH protein in *Vhl* KO kidneys confirmed by Western Blot (1.00 ± 0.11 vs 0.32 ± 0.079). Relative protein expression is normalized to total protein. **E.** DNA dot blot for 5-hmC in *Vhl* WT and *Vhl* KO kidneys. Methylene blue shown as loading control. Relative 5-hmC abundance is normalized to methylene blue loading control.

### Belzutifan reverses miRNA target gene suppression and increases 5-hmC

Belzutifan is a potent small-molecule HIF2α inhibitor recently approved in the treatment of VHL disease associated tumors, and also in sporadic advanced ccRCC. We sought to determine the efficacy of belzutifan in targeting miR155-5p and miR21-3p in ccRCC cell lines (**Figure 7**). In A-704 cells, HIF2α inhibition was confirmed by decreased expression of HIF2α targets *VEGFA* and *CXCR4* (**Figure 7A**). Also, treatment with belzutifan resulted in inhibition of both miR21-3p and miR155-5p in A-704 cells (**Figure 7A**), with a concomitant increase in *L2HGDH* and *HIF1A* target gene expression (**Figure 7A**). This was replicated in the 786-O cell line with confirmation of HIF2α inhibition, decreased miRNA expression, and increased *L2HGDH* and *HIF1A* gene expression (**Figure 7B**). We then measured 5-hmC to determine whether the increase in *L2HGDH* would be associated with restoration of 5-hmC levels. Indeed, belzutifan treatment resulted in significant increases in 5-hmC content in both A-704 and 786-O cell lines (**Figure 7C, 7D**).

**Figure 7.**
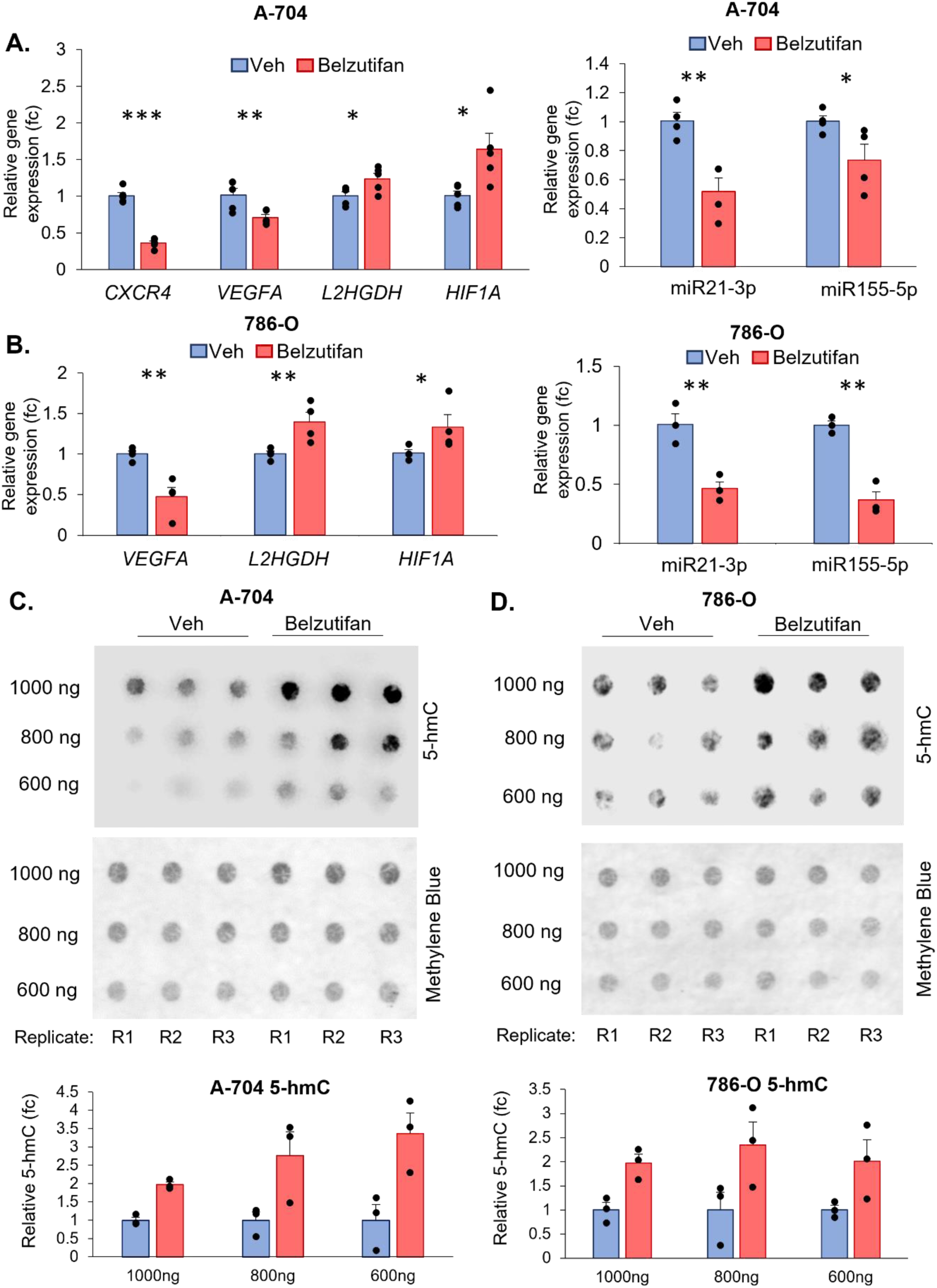
HIF2α inhibition with belzutifan reverses miR21-3p and miR155-5p target gene suppression and increases 5-hmC in ccRCC cells. **A.** A-704 cells were treated with 2 µM belzutifan or Veh for 24 h. HIF2α inhibition was confirmed by downregulation of HIF2α target genes *VEGFA* (n=5, P=0.0075) and *CXCR4* (n=5, P<0.001). Belzutifan treatment resulted in upregulation of *L2HGDH* (n=5, P=0.020) and *HIF1A* (n=5, P=0.013). Belzutifan also decreased miR21-3p (n=4, P=0.0023) and miR155-5p (n=4, P=0.030) expression. [P-values represent one-sided t-tests. *, p<0.05; **, p<0.01; ***, p<0.001] **B.** 786-O cells were treated with 20 µM belzutifan or vehicle (Veh) for 12 h (n=3 per condition). Belzutifan treatment resulted in upregulation of *L2HGDH* (P= 0.0097) and *HIF1A* (P=0.044) as well as downregulation of HIF2α target gene *VEGFA* (P=0.0025). Belzutifan treatment also resulted in decreased miR21-3p (P=0.0049) and significant inhibition of miR155-5p (P=0.0010) expression. [P-values represent one-sided t-tests. *, p<0.05; **, p<0.01; ***, p<0.001] **C, D**. DNA 5-hmC dot blots showed a robust increase in 5-hmC content with belzutifan treatment in both A-704 and 786-O cell lines.

### Combined inhibition of HIF1α and HIF2α in ccRCC cells expressing both isoforms

Because HIF1α and HIF2α partially overlap in their DNA binding sites, we sought to determine the relative contribution of each to regulation of *MIR21* and *MIR155*, and whether dual inhibition of HIF1α and HIF2α may result in greater inhibition of target genes. For these experiments, we used the RCC4 cell line which, unlike 786-O and A-704, has intact *HIF1A* transcript. ChIP-seq for HIF1α and HIF2α in the RCC4 cell line showed that both HIF1α and HIF2α bind to the *MIR155* promoter (**Figure 8A**). However, unlike 786-O and A-704, HIF2α showed only minimal enrichment at the *MIR21* site, with no peak observed for HIF1α binding (**Figure 8B**). We then compared knockdown of *HIF1A* and *EPAS1*, individually and in combination, on miRNA and target gene expression. Knockdown was confirmed for each gene at the transcript level, and respective downregulation of HIF1α-specific (*CA9*) and HIF2α -specific (*VEGFA*) target genes was also confirmed (**Fig 8C**). Knockdown of both *HIF1A* and *EPAS1* resulted in a decrease in miR155-5p expression, with the most robust decrease in the *EPAS1* knockdown condition (**Figure 8D**). Consistent with this, we also observed increased expression of the miR155-5p target gene, *HIF1A*, in the *EPAS1* knockdown condition (**Fig 8C**). However, in line with ChIP-seq data, miR21-3p expression was found to be low in this cell line, limiting the use of this model to study miR21-3p regulation. As such, knockdown of *HIF1A* and *EPAS1* did not significantly decrease expression of miR21-3p further (**Figure 8D**). However, the miR21-3p target gene, *L2HGDH*, was increased by *HIF1A* knockdown, and to a greater extent by *EPAS1* and combined *HIF1A*/*EPAS1* knockdown. Given the low expression of miR21-3p in RCC4, there are likely additional mechanisms resulting in HIF-induced repression of *L2HGDH* in this cell line. Given the observed increase in *L2HGDH*, we assessed 5-hmC in these conditions.

**Figure 8.**
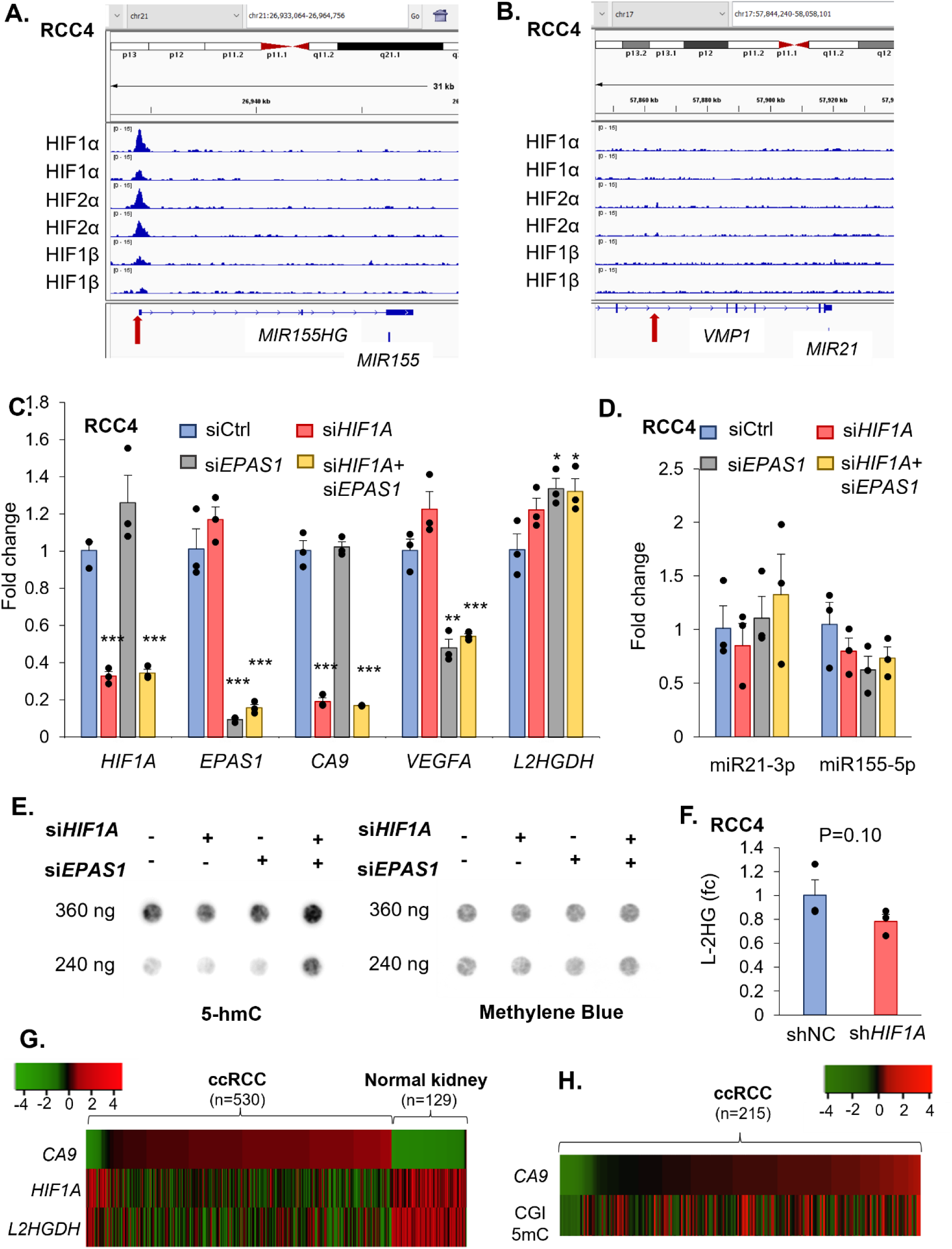
Simultaneous inhibition of HIF2α and HIF1α in ccRCC cells with upregulation of both isoforms. **A, B.** Visualization of HIF1α, HIF2α, and HIF1β (n=2 each) ChIP-seq data for the RCC4 cell line (GSE120885^19^). Binding peaks are shown for (**A**) *MIR155* and (**B**) *MIR21* loci. **C, D**. siRNA targeting control, *HIF1A*, *EPAS1*, and *HIF1A*/ *EPAS1* were transfected in the RCC4 cell line. Relative gene expression was determined 48-h post-transfection (n=3 per transfection). **C.** *HIF1A* and *EPAS1* gene expression confirmed knockdown of respective transcripts. *CA9* and *VEGFA* further confirmed knockdown of HIF1α and HIF2α target genes, respectively. *L2HGDH* was elevated slightly by *HIF1A* knockdown, and to a greater extent by *EPAS1* and *HIF1A*/ *EPAS1* knockdown. **D.** miR21-3p has low expression in RCC4 and is unchanged by *HIF1A* and *EPAS1* knockdown. miR155-5p is slightly decreased by knockdown of HIF1A (P>0.1) and to a greater extent by *EPAS1* knockdown (P=0.078). [ *, p<0.05; **, p<0.01; ***, p<0.001; one-sided t-test] **E**. DNA dot blot showing 5-hmC in the RCC4 cell line 72-h post-transfection with *HIF1A* and *EPAS1* targeting siRNA. 5-hmC is increased robustly with combined *HIF1A* and *EPAS1* knockdown. **F**. L-2HG analysis of RCC4 cell line following *HIF1A* shRNA stable knockdown (n=3) compared to non-targeting shRNA control (shNC, n=3). L-2HG is decreased by *HIF1A* knockdown (0.78 ± 0.062) compared to control (1.0 ± 0.13). Data are shown as mean ± SE. Knockdown verification and D-2HG are shown in Suppl Fig S7. **G.** Heatmap of *CA9*, *HIF1A*, *L2HGDH* expression in ccRCC (TCGA KIRC) demonstrating a marked negative correlation between *CA9* expression (marker of HIF1α transcriptional activity) and *HIF1A* expression (R= -0.55; p=1.8e-52), as well as a similarly marked negative correlation between *CA9* and *L2HGDH* expression (R= -0.59; p=9.3e-64). **H.** Heatmap demonstrating a significant positive correlation between *CA9* expression and CpG island methylation in ccRCC (TCGA KIRC) (R=+0.38, p=8.6e-9)

Combined knockdown of *HIF1A* and *EPAS1* resulted in a gain of 5-hmC, but knockdown of either alone did not (**Fig 8E**). [We have previously established that loss of 5-hmC is an independent adverse prognostic feature of ccRCC. This loss of 5-hmC in ccRCC is not only secondary to under-expression of L2HGDH (and subsequent accumulation of L2HG) but also under-expression of SDH (and subsequent succinate accumulation.)].^1, 30^ Finally, to explore the effect of *HIF1A* knockdown on L-2HG abundance, we assessed L-2HG by LC-MS in RCC4 stable shRNA *HIF1A* knockdown cells (**Fig 8F, Suppl Fig S7A**). Knockdown of *HIF1A* resulted in 21.8% reduction in L-2HG compared to non-targeting control shRNA cells (**Fig 8F**). D-2HG was not affected by *HIF1A* knockdown (**Suppl Fig S7B**).

Consistent with in vitro findings, TCGA KIRC data demonstrated a marked negative correlation between *CA9* expression (marker of HIF1α transcriptional activity) and *HIF1A* expression (R= - 0.55; P = 1.8e-52), as well as a similarly marked negative correlation between *CA9* and *L2HGDH* expression (R = -0.59; P = 9.3e-64, **Figure 8G**). Furthermore, there was significant positive correlation between *CA9* expression and CpG island methylation (R= +0.38, P = 8.6e-9, **Figure 8H**).

To further confirm the regulatory relationship, we treated the normal kidney cell line, HKC-8, with the hypoxia mimetic agent/ prolyl hydroxylase inhibitor, dimethyloxalylglycine (DMOG) and measured RNA and protein expression of HIFα, miRNAs, and target genes (**Fig 9**). Consistent with the hypothesis, despite the characteristic increase in HIF2α and HIF1α protein expression with DMOG treatment, *HIF1A* transcript levels decreased, while *HIF2A* transcript levels increased (**Fig 9A, 9B**), further demonstrating the negative feedback for *HIF1A* but not *HIF2A*. L2HGDH decreased both at the transcript and protein levels with DMOG treatment (**Fig 9A, 9B**) We also observed a significant increase in miR155-5p and miR21-3p expression with DMOG treatment, as hypothesized (**Fig 9C**). Together, these data support the conserved inhibitory feedback mechanism for HIFα/ *HIF1A* as well as RNA transcript inhibition of miRNA target genes (↑HIFα→↑miR155-5p→↓*HIF1A;* ↑HIFα→↑miR21-3p→↓*L2HGDH*).

**Figure 9.**
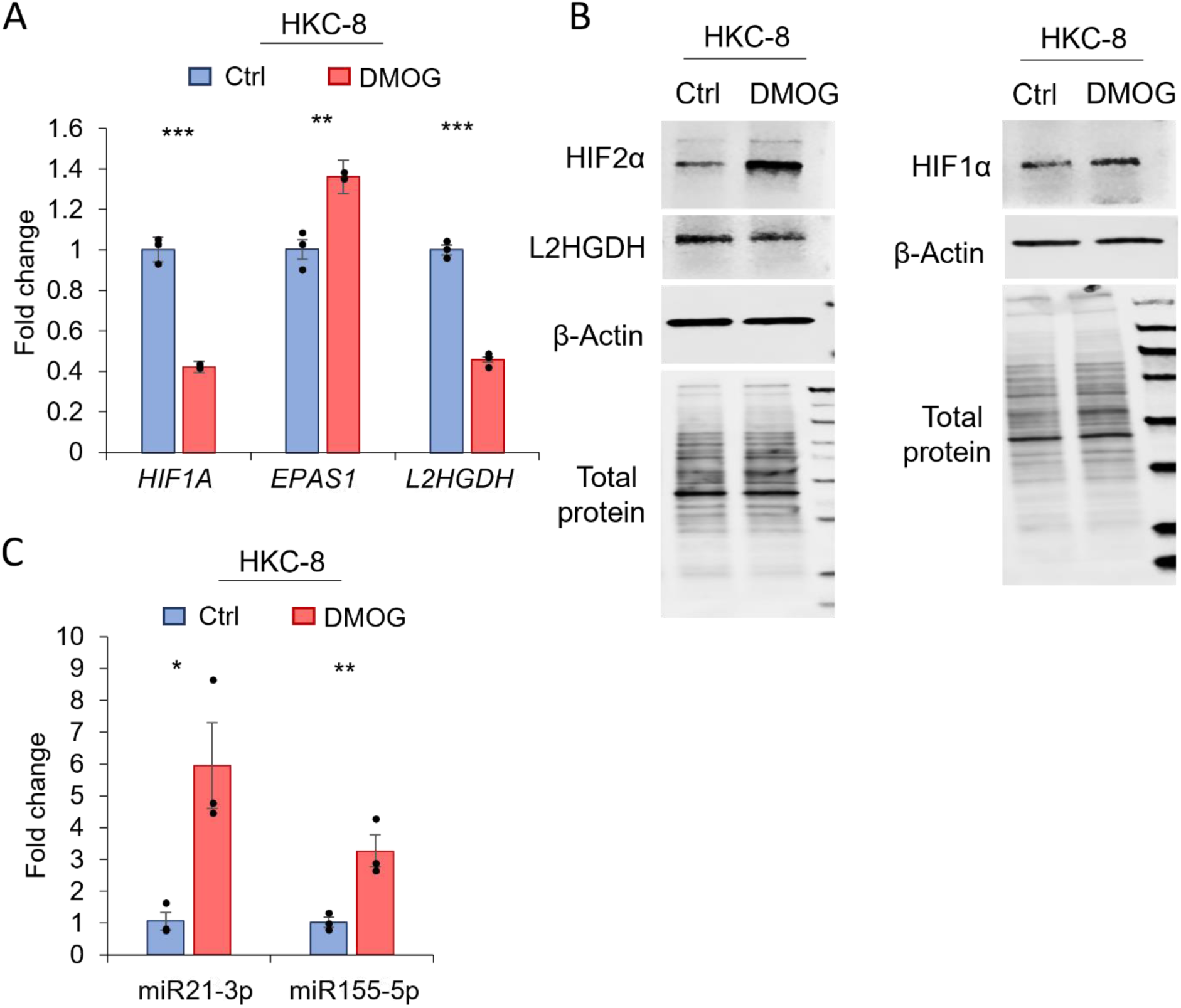
Hypoxia mimetic agent dimethyloxalylglycine (DMOG) induces the conserved inhibitory HIFα/ *HIF1A* feedback loop and downregulates L2HGDH in normal kidney cells. **A-B.** HKC8 cells were exposed to 1 mM DMOG. Protein and gene expression were assessed by qPCR (**A,C**) and Western Blot (**B**). DMOG treatment resulted in decreased *HIF1A* and increased *HIF2A* transcript levels (**A**), along with the expected increase in HIF2α and HIF1α protein (**B).** DMOG treatment increased miR21-3p and miR155-5p expression (**C**); and decreased L2HGDH both at the transcript and protein levels (**A,B**). [ *, p<0.05; **, p<0.01; ***, p<0.001; one-sided t-test]

## Discussion

In this study, we sought to identify mechanisms governing the downregulation of *L2HGDH* and *HIF1A* in ccRCC independent of 14q deletion. While lower *L2HGDH* portends strikingly worse prognosis in a multivariate analysis (Figure 1B) highlighting its translational importance, the lower expression of *HIF1A* in ccRCC (contrary to the upregulation to *HIF2A*) is of significant scientific interest.

We show that the downregulation of both genes in ccRCC is mediated by microRNAs that are markedly elevated compared to normal kidney, secondary to pseudo hypoxic signaling. Using bioinformatic analyses of large clinicogenomic and proteomic datasets (TCGA KIRC and CPTAC), identification of baseline ccRCC and pre-cancerous cell line expression of relevant genes/ miRNAs and changes with genetic manipulation (over-expression/ knockdown), reporter assay for specificity of miRNA targets, chromatin immunoprecipitation and sequencing analyses, we show that the hypoxia inducible factors upregulate miR21-3p and miR155-5p, which subsequently target *L2HGDH* and *HIF1A* respectively. Using a mouse model of kidney specific *Vhl* deletion, we show that loss of *Vhl* and consequent pseudohypoxic signaling is sufficient to upregulate miR155-5p and miR21-3p, down-regulate *L2hgdh* resulting in loss of hydroxymethylcytosine. Finally, we demonstrate that inhibition of HIF can reverse these metabolic and epigenetic aberrations.

### A conserved inhibitory feedback loop for *HIF1A,* but not *EPAS1,* in ccRCC

Despite the high expression of HIF1α and HIF2α at the protein level secondary to VHL loss, *HIF1A* is under-expressed compared to normal kidney (but *EPAS1,* encoding HIF2α, is over-expressed). In this study, we show that only *HIF1A* but not *EPAS1* is targeted by HIF-induced miR155-5p for degradation, providing an explanation for the differential expression at the transcript level. This is likely an evolutionarily conserved mechanism to regulate responses to hypoxia in normal cells. We show that in response to a hypoxia mimetic agent, normal (immortalized) kidney cells upregulate HIFα proteins, upregulate miR155-5p, and downregulate *HIF1A* (but not *EPAS1*). Of note, miR155-5p has been shown to target *HIF1A* in intestinal epithelial cells as well.^31^ The promoters of both *EPAS1* and *HIF1A* show binding sites for HIF1α and HIF2α similar to many transcription factors’ positive self-regulation, but this results in a positive upregulation of only *EPAS1* expression but not *HIF1A* due to the latter’s miRNA degradation (via miR155-5p and potentially other miRNAs). However, given that the primary mode of regulation of HIFα isoforms is post-translational, the under expression of *HIF1A* appears to have little effect on the protein expression of HIF1α in ccRCC with VHL loss, and the degree of protein upregulation of HIF1α is similar to that of HIF2α in VHL-deficient ccRCC.^8^ (also see Figures S6A-D). But in normal VHL intact cells, this inhibitory feedback loop to lower *HIF1A* plausibly influences HIF1α levels to regulate the degree of chronic hypoxic response.

### HIF inhibition reverses metabolic and epigenetic aberrations in ccRCC

Our study demonstrates that HIFs drive early and sustained metabolic and epigenetic dysregulation in ccRCC. These HIF-driven aberrations begin early in carcinogenesis. We show that inhibition of HIFs can reverse these pathological changes - most significantly, restoring levels of 5-hydroxymethylcytosine, a key epigenetic marker lost in ccRCC due to metabolic inhibition of TET enzymes. Our findings support the integration of HIF2α inhibition earlier in the therapeutic approach for advanced ccRCC.

In ccRCC cells which only express/ upregulate the HIF2α isoform (786-O and A-704), selective pharmacologic inhibition of HIF2α with belzutifan led to downregulation of miR21-3p, upregulation of L2HGDH, and gain of hydroxymethylcytosine. These data indicate that in ccRCC tumors with high-HIF2/ low-HIF1, belzutifan can (at least partially) reverse the metabolic and epigenetic aberrancies in ccRCC, However, in ccRCC cells with upregulation of both HIF1α and HIF2α isoforms (RCC4), knockdown of either *EPAS1* or *HIF1A* alone did not reverse the loss of hydroxymethylcytosine, but combined inhibition did. These data indicate that in ccRCC tumors with high-HIF2α / high-HIF1α or low-HIF2α /high-HIF1α, only inhibition of HIF2α is unlikely to significantly reverse the adverse epigenetic aberrations, but combined HIF2α and HIF1α inhibition can. This warrants further investigation in future studies given its strong translational significance. While there may be concern regarding the potential toxicity of dual HIF targeting, there are non-toxic agents capable of modulating HIF1α activity - either by inhibiting transcriptional activity or promoting degradation - that could be investigated in future combination studies.

The LITESPARK-003 trial, which is evaluating belzutifan in combination with cabozantinib (a tyrosine kinase inhibitor) in previously untreated advanced ccRCC patients, has recently reported highly promising results, with an overall response rate of 70%. It would be both interesting and important to correlate HIF2α/ HIF1α status with treatment response, and to assess whether non-responders exhibited higher HIF1α levels.

In summary, this study sheds further light on the intricate links between constitutionally hyperactivated hypoxia signaling, metabolic dysregulation and epigenetic aberrancies in clear cell renal cancer, during pathogenesis and progression. It reveals that HIFs drive key metabolic and epigenetic aberrancies in clear cell renal cancer, and that inhibiting HIFs can at least partially reverse the aberrancies. It also unveils a conserved inhibitory feedback loop for *HIF1A* in ccRCC, adding to the intriguing dynamics between the HIFα isoforms in this malignancy.

## Data sharing statement

Data sources and software used are specified throughout.

## Acknowledgments

Sources of support: This work was supported by funding from the American Cancer Society Research Scholar Grant RSG-20-137-01 TBG (NKS), the Northwestern University Division of Hematology/ Oncology (NKS), the Lotte & John Hecht Foundation grant #4809 (NKS), Roswell Park Comprehensive Cancer Center and National Cancer Institute (NCI) grant, P30CA016056. Metabolomics experiments (L2HG) were performed at the Metabolomics Core Facility at Robert H. Lurie Comprehensive Cancer Center (supported by NCI CCSG P30 CA060553) of Northwestern University.

## Author Contributions

SS, RAL and NKS designed the experiments; SS, RAL, TA performed the experiments; SS, RAL, TA, and NKS analyzed the data; NKS performed the bioinformatic analyses; RAL and NKS wrote the manuscript; BN, BI, KP, SH edited the manuscript, NKS was responsible for conceptualization, direction, and funding.

## List of supplementary material

-Supplementary Figure Document

-Full uncropped blots

## SUPPLEMENTAL MATERIAL/ EXTENDED DATA

**Fig S1.**
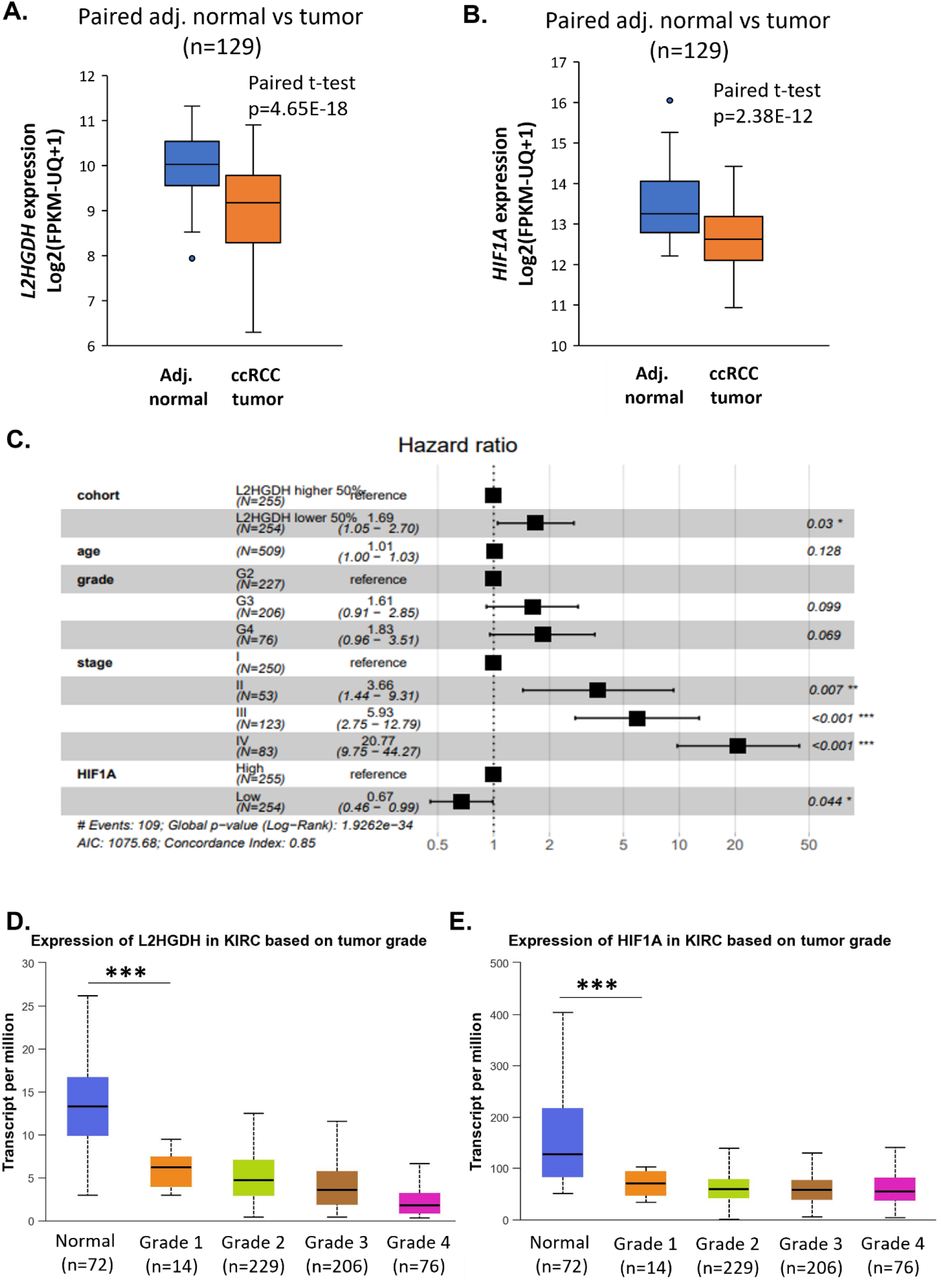
TCGA KIRC data revealing significant downregulation of both *L2HGDH* (**A**) and HIF1A (**B**) in ccRCC compared to paired adjacent normal kidney. (**C.**) Cox regression multivariate survival analysis with TCGA KIRC reveals that the lower half of *L2HGDH* expression is associated with a 69% increase in hazard of death from ccRCC compared to the upper half, after adjusting for grade, stage, age, and *HIF1A* expression, (HR 1.69; 95% CI: 1.05-2.70; p=0.03). Expression of *L2HGDH* (**D**) and *HIF1A* (**E**) in ccRCC revealing a marked downregulation already in grade 1 disease, compared to normal kidney (TCGA KIRC).

**Fig S2.**
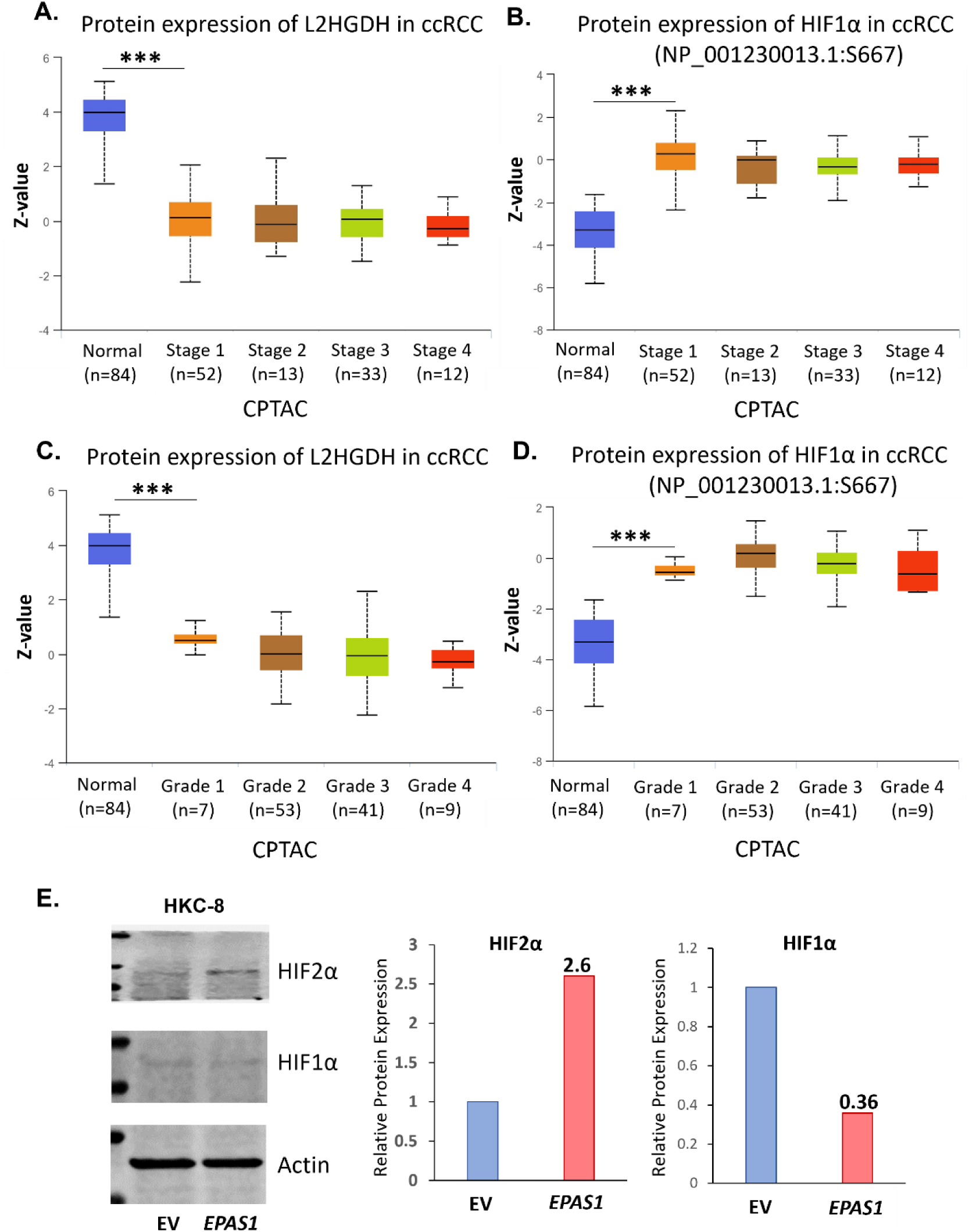
Protein expression of L2HGDH (**A, C**) and phospho-protein expression of HIF1α (**B, D)** stratified by Stage (**A, B**) and Grade (**C, D**) in the CPTAC dataset, revealing a marked downregulation of both early in disease progression **E**. Expression of HIF2α and HIF1α following *EPAS1* overexpression in HKC-8 cell line.

**Fig S3.**
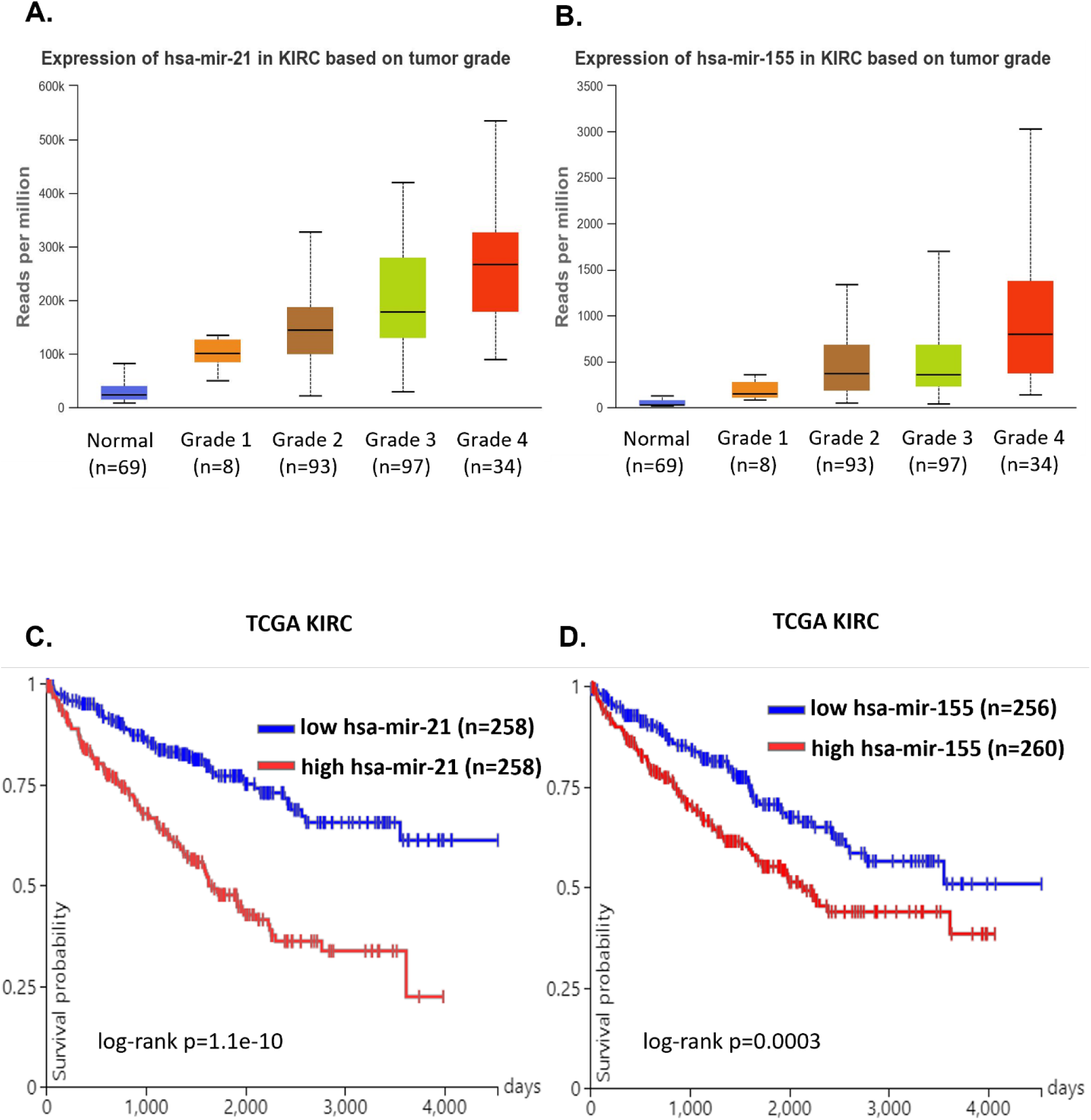
**A, B**. Expression of miR21 and miR155 in normal kidney and ccRCC tumors stratified by Grade. **C, D.** Survival based on stratification by miR21 and miR155 expression (Higher expression of both miR21 and miR155 is associated with significantly worse survival in a univariate analysis). All samples were part of the TCGA KIRC.

**Fig S4.**
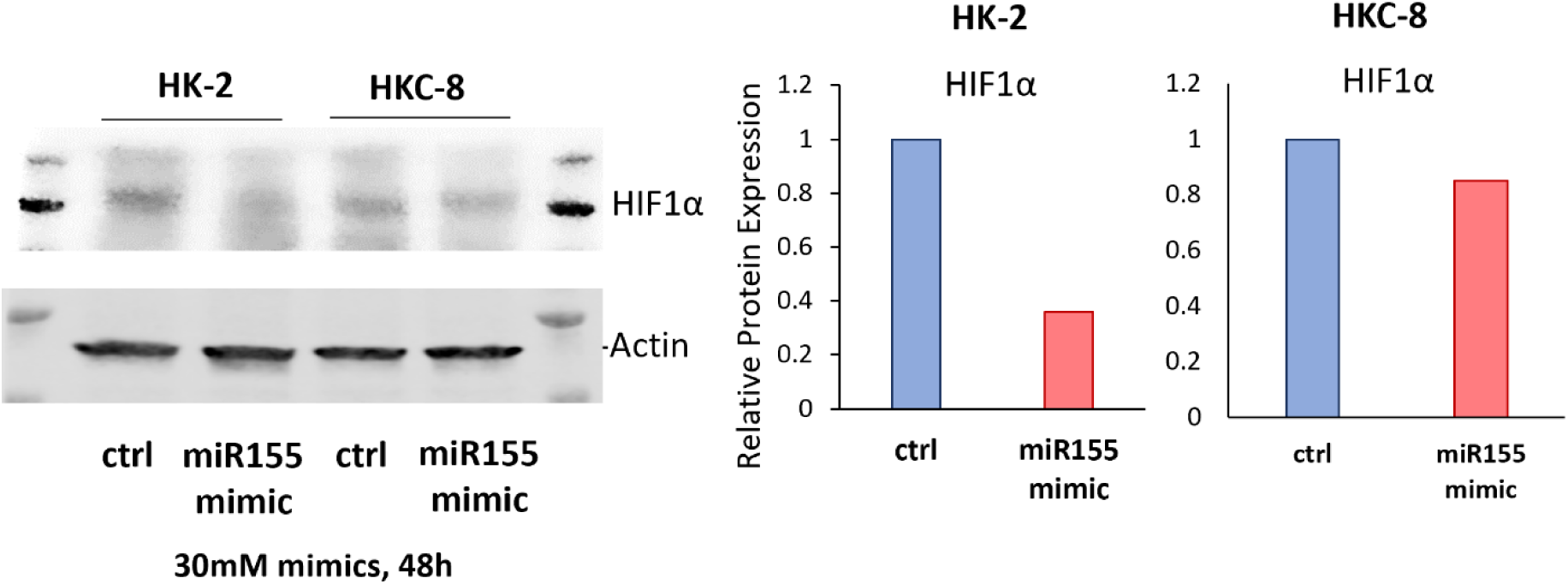
HIF1α protein expression in HK-2 and HKC-8 cell lines transfected with non-targeting control of miR155 siRNA for 48 hours.

**Fig S5.**
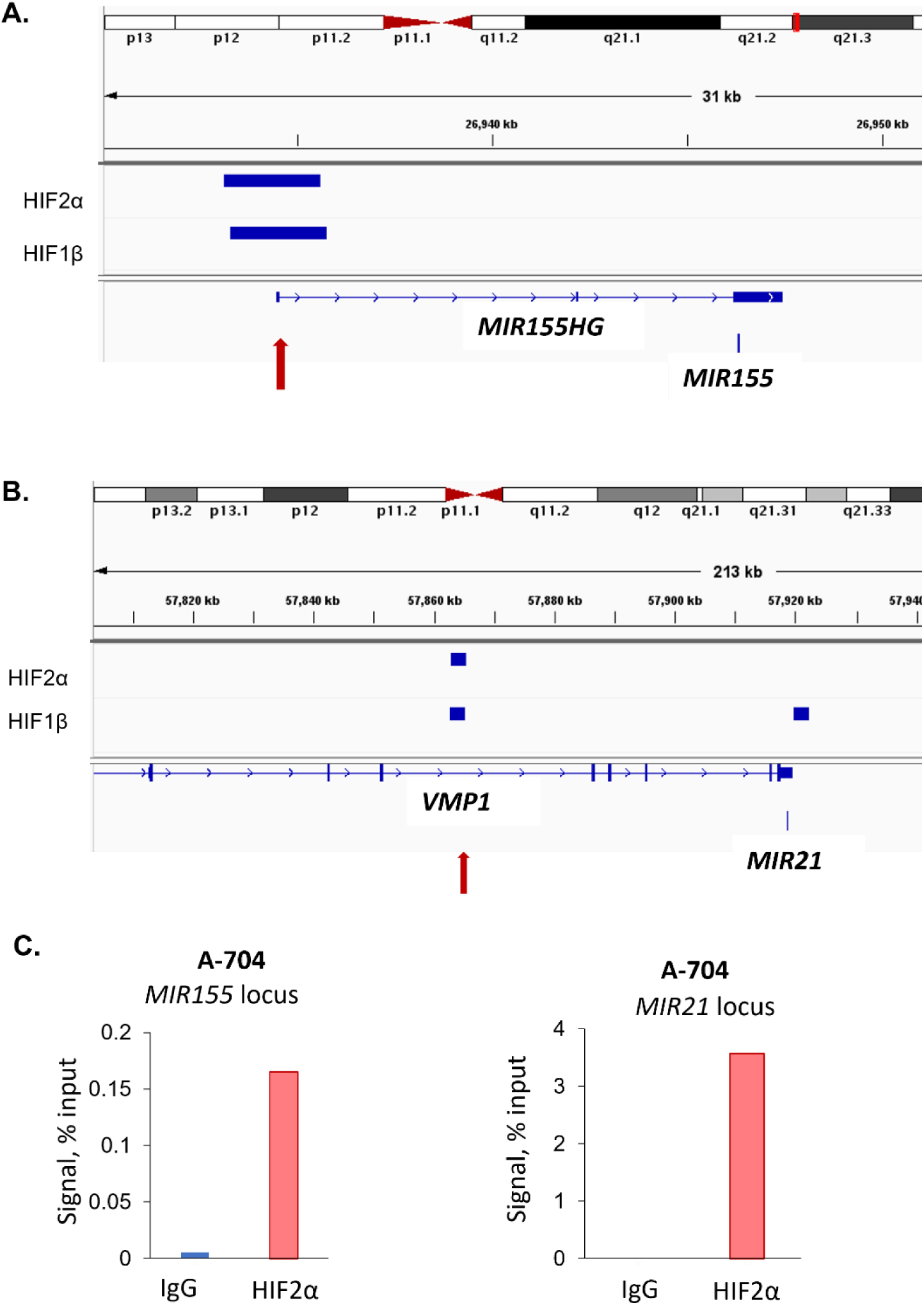
**A,B.** ChIP for HIF2α and HIF1β in 786-O cells. ChIP-seq was visualized from GSE67237 (pmid 26262842). C. HIF2α ChIP in A-704 cell line confirming HIF2α binding at both *MIR155* and *MIR21* loci.

**Fig S6.**
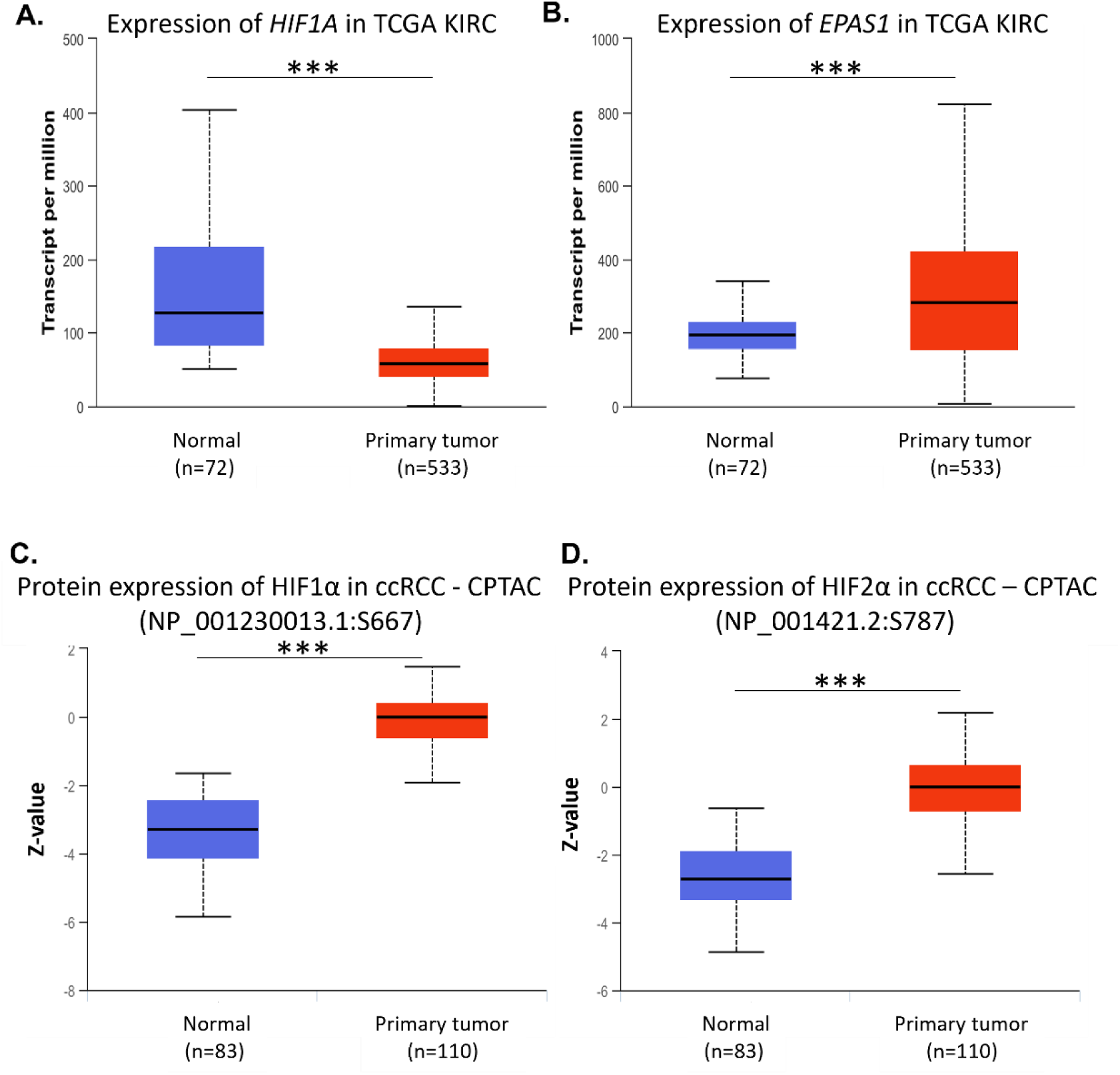
**A, B.** Gene expression of *HIF1A* and *EPAS1* in TCGA KIRC. **C.D.** Phospho-protein expression of HIF1α and HIF2α in CPTAC.

**Fig S7.**
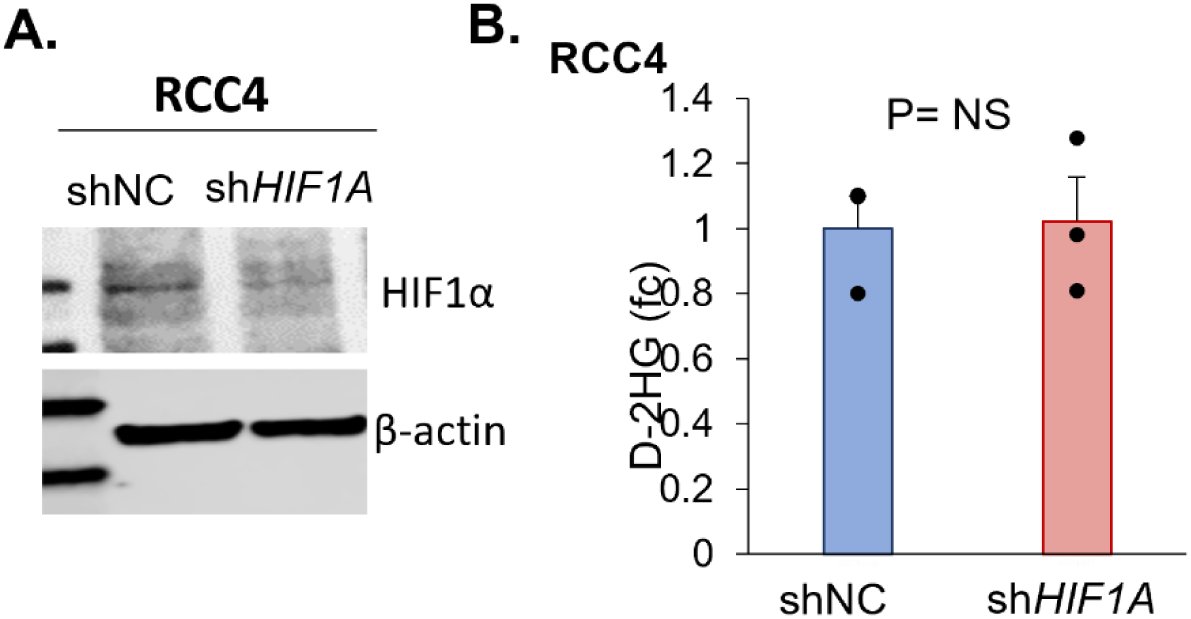
**A.** *HIF1A* shRNA knockdown validation by Western blot in RCC4 cells. **B.** D-2HG is not different between shNC (non-targeting control) and sh*HIF1A* RCC4 cells.

## References

1. Shenoy N, Bhagat TD, Cheville J, et al. Ascorbic acid-induced TET activation mitigates adverse hydroxymethylcytosine loss in renal cell carcinoma. J Clin Invest 2019; 130: 1612–1625.

2. Shenoy N, Pagliaro L. Sequential pathogenesis of metastatic VHL mutant clear cell renal cell carcinoma: putting it together with a translational perspective. Ann Oncol 2016; 27: 1685–1695.

3. Hakimi AA, Reznik E, Lee CH, et al. An Integrated Metabolic Atlas of Clear Cell Renal Cell Carcinoma. Cancer Cell 2016; 29: 104–116.

4. Monzon FA, Alvarez K, Peterson L, et al. Chromosome 14q loss defines a molecular subtype of clear-cell renal cell carcinoma associated with poor prognosis. Mod Pathol 2011; 24: 1470–1479.

5. Shenoy N. HIF1α is not a target of 14q deletion in clear cell renal cancer. Sci Rep 2020; 10: 17642.

6. Shim EH, Livi CB, Rakheja D, et al. L-2-Hydroxyglutarate: an epigenetic modifier and putative oncometabolite in renal cancer. Cancer Discov 2014; 4: 1290–1298.

7. Shelar S, Shim EH, Brinkley GJ, et al. Biochemical and Epigenetic Insights into L-2-Hydroxyglutarate, a Potential Therapeutic Target in Renal Cancer. Clin Cancer Res 2018; 24: 6433–6446.

8. Krieg M, Haas R, Brauch H, et al. Up-regulation of hypoxia-inducible factors HIF-1alpha and HIF-2alpha under normoxic conditions in renal carcinoma cells by von Hippel-Lindau tumor suppressor gene loss of function. Oncogene 2000; 19: 5435–5443.

9. Di Cristofano C, Minervini A, Menicagli M, et al. Nuclear expression of hypoxia-inducible factor-1alpha in clear cell renal cell carcinoma is involved in tumor progression. Am J Surg Pathol 2007; 31: 1875–1881.

10. Klatte T, Seligson DB, Riggs SB, et al. Hypoxia-inducible factor 1 alpha in clear cell renal cell carcinoma. Clin Cancer Res 2007; 13: 7388–7393.

11. Minardi D, Lucarini G, Filosa A, et al. Prognostic role of tumor necrosis, microvessel density, vascular endothelial growth factor and hypoxia inducible factor-1alpha in patients with clear cell renal carcinoma after radical nephrectomy in a long term follow-up. Int J Immunopathol Pharmacol 2008; 21: 447–455.

12. Shen C, Beroukhim R, Schumacher SE, et al. Genetic and functional studies implicate HIF1α as a 14q kidney cancer suppressor gene. Cancer Discov 2011; 1: 222–235.

13. Fu L, Wang G, Shevchuk MM, et al. Generation of a mouse model of Von Hippel-Lindau kidney disease leading to renal cancers by expression of a constitutively active mutant of HIF1α. Cancer Res 2011; 71: 6848–6856.

14. Hoefflin R, Harlander S, Schäfer S, et al. HIF-1α and HIF-2α differently regulate tumour development and inflammation of clear cell renal cell carcinoma in mice. Nat Commun 2020; 11: 4111.

15. Cowman SJ, Koh MY. Revisiting the HIF switch in the tumor and its immune microenvironment. Trends Cancer 2022; 8: 28–42.

16. Kondo K, Klco J, Nakamura E, et al. Inhibition of HIF is necessary for tumor suppression by the von Hippel-Lindau protein. Cancer Cell 2002; 1: 237–246.

17. Schodel J, Bardella C, Sciesielski LK, et al. Common genetic variants at the 11q13.3 renal cancer susceptibility locus influence binding of HIF to an enhancer of cyclin D1 expression. Nat Genet 2012; 44: 420–425, s421-422.

18. Salama R, Masson N, Simpson P, et al. Heterogeneous Effects of Direct Hypoxia Pathway Activation in Kidney Cancer. PLoS One 2015; 10: e0134645.

19. Smythies JA, Sun M, Masson N, et al. Inherent DNA-binding specificities of the HIF-1α and HIF-2α transcription factors in chromatin. EMBO Rep 2019; 20.

20. Wang J, Dai X, Berry LD, et al. HACER: an atlas of human active enhancers to interpret regulatory variants. Nucleic Acids Res 2019; 47: D106–D112.

21. Percie du Sert N, Hurst V, Ahluwalia A, et al. The ARRIVE guidelines 2.0: Updated guidelines for reporting animal research. PLoS Biol 2020; 18: e3000410.

22. Goldman MJ, Craft B, Hastie M, et al. Visualizing and interpreting cancer genomics data via the Xena platform. Nat Biotechnol 2020; 38: 675–678.

23. Cerami E, Gao J, Dogrusoz U, et al. The cBio cancer genomics portal: an open platform for exploring multidimensional cancer genomics data. Cancer Discov 2012; 2: 401–404.

24. Guan X, Cai M, Du Y, et al. CVCDAP: an integrated platform for molecular and clinical analysis of cancer virtual cohorts. Nucleic Acids Res 2020; 48: W463–w471.

25. Chandrashekar DS, Bashel B, Balasubramanya SAH, et al. UALCAN: A Portal for Facilitating Tumor Subgroup Gene Expression and Survival Analyses. Neoplasia 2017; 19: 649–658.

26. Robinson JT, Thorvaldsdottir H, Winckler W, et al. Integrative genomics viewer. Nat Biotechnol 2011; 29: 24–26.

27. Nassar LR, Barber GP, Benet-Pagès A, et al. The UCSC Genome Browser database: 2023 update. Nucleic Acids Res 2023; 51: D1188–d1195.

28. Chen Y, Wang X. miRDB: an online database for prediction of functional microRNA targets. Nucleic Acids Res 2020; 48: D127–d131.

29. Wong NW, Chen Y, Chen S, et al. OncomiR: an online resource for exploring pan-cancer microRNA dysregulation. Bioinformatics 2018; 34: 713–715.

30. Aggarwal RK, Luchtel RA, Machha V, et al. Functional succinate dehydrogenase deficiency is a common adverse feature of clear cell renal cancer. Proc Natl Acad Sci U S A 2021; 118.

31. Bruning U, Cerone L, Neufeld Z, et al. MicroRNA-155 promotes resolution of hypoxia-inducible factor 1alpha activity during prolonged hypoxia. Mol Cell Biol 2011; 31: 4087–4096.

32. Schönenberger D, Rajski M, Harlander S, et al. Vhl deletion in renal epithelia causes HIF-1α-dependent, HIF-2α-independent angiogenesis and constitutive diuresis. Oncotarget 2016; 7: 60971–60985.

